# Social experience alters behaviors by reprogramming the Fruitless pathway and circadian state in *Drosophila*

**DOI:** 10.1101/2025.05.30.657129

**Authors:** Chengcheng Du, Sumie Okuwa, Lanling Jia, Marta Rozados Barreiro, Shayna Scott, Jesús Emiliano Sotelo Fonseca, Yuta Mabuchi, Shania Appadoo, Luis Garcia, Stephanie Rohrbach, Sare Koruk, Efe Balkanli, Josh Lin, Nilay Yapici, Corbin D. Jones, Pelin C. Volkan

## Abstract

From flies to humans, social experience affects various cognitive and behavioral processes. Previous studies have shown that group housing suppresses many behaviors like courtship, aggression, and feeding in *Drosophila melanogaster,* in addition to resetting the circadian state. Here, we focus on group housing-induced courtship suppression. To determine the mechanisms by which social experience modulates courtship behaviors, we performed bulk tissue RNAseq and single-cell RNAseq from cells expressing Fruitless^M^ (Fru^M^) and Doublesex^M^ (Dsx^M^), two transcription factors that label interconnected neural circuits for socially driven behaviors, from grouped or isolated male brains. These revealed that social isolation alters *fru* and *dsx* levels throughout the brain. Knocking down *fru^M^* in different *fru^M^*-positive neuron subpopulations in the brains has diverse effects on social experience-dependent changes in courtship. Furthermore, group housing increases the expression of *stripe* (*sr*) and *Hormone receptor-like in 38* (*Hr38*) genes encoding neural activity-induced transcription factors in most neurons within social circuits. We found that knocking down *sr* in *fru^M^*-positive neurons effectively eliminates the impact of social experience by increasing courtship in group-housed males. Importantly, social experience also alters the expression of Fru^M^/Dsx^M^ putative target genes regulating circadian states throughout the brain. Disrupting the function of multiple circadian genes diminishes the effect of group housing on courtship. Our findings suggest that group housing/social enrichment suppresses courtship by reprogramming the circadian arousal state, whereas courtship-elevating effects of social experience rely on unique influences of Fru^M^ expression and function in different neurons within social and clock circuits. These results are significant as they point to modulation of circadian arousal state as a possible central strategy for mediating the pleiotropic effects of social experience on organismal responses.

## Introduction

Social experience, particularly social isolation or loneliness, is known to have long-lasting effects on behavior and physiology in animals ranging from mice to flies (Anders, 1978; Cacioppo et al., 2006; de Melo Sene, 1977; Hoffman et al., 1975; Sánchez et al., 1998). Over the past decade, growing awareness of these effects—exacerbated by global health crises—has led to the recognition of social isolation and loneliness as a new epidemic, as declared by the Surgeon General (General, 2023; Jeste et al., 2020; Lin, 2023). Adverse social experience, including isolation, disrupts various cognitive processes, including sleep, learning, memory, and attention, while also increasing behaviors such as aggression and feeding (Brown et al., 2017; Cacioppo and Hawkley, 2009; Cacioppo et al., 2006; Cohen, 2004; Liu et al., 2025; Zelikowsky et al., 2018). Consequently, in humans, isolation is linked to a rise in neuropsychiatric and neurodegenerative diseases (Cacioppo et al., 2015; Cacioppo et al., 2006; Cohen, 2004). The molecular and circuit-based mechanisms of social experience-dependent behavioral and physiological effects remain unclear.

Social experience similarly shapes physiology and behavior across animal species. Animals modulate their behaviors by signals from the social environment and prior social experience (Anders, 1978; de Melo Sene, 1977; Eban-Rothschild and Bloch, 2015; Luciano and Lore, 1975; Matsumoto et al., 2005; Ueda and Kidokoro, 2002; Wang et al., 2022). Emerging evidence from both vertebrate and invertebrate models demonstrates that social environment exerts profound effects on the regulation of genes critical for maintaining neuronal homeostasis, physiological function, and synaptic transmission, ultimately shaping neural circuit activity and behavioral outputs (Cushing and Kramer, 2005; Deanhardt et al., 2023; Flavell and Greenberg, 2008; West and Greenberg, 2011; Zhao et al., 2020). Despite this progress, how the detection of social cues by peripheral sensory systems alters gene expression throughout central circuits to ultimately modify circuit function and behaviors remains poorly understood.

Exploring molecular and circuit mechanisms by which social experience modulates behaviors requires a well-defined model system with causal links between genes, circuits, and behavioral function. The genetically tractable fruit fly is an excellent model because of its adaptable social behaviors, simple nervous system with a widely mapped connectome, and known master regulatory genes for circuit structure and function. Beyond the general advantages of *Drosophila* as a model organism, its pronounced sensitivity to social experience makes it particularly valuable for identifying genes and circuits driving responses to social experience. Previous studies reported that social isolation elevates many fundamental behaviors, like feeding, aggression, and courtship, while disrupting cognitive processes such as sleep, learning, and memory in fruit flies (Agrawal et al., 2020; Dankert et al., 2009; Ganguly-Fitzgerald et al., 2006; Goncharova et al., 2016; Kacsoh et al., 2018; Li et al., 2021; Ueda and Kidokoro, 2002; Wang et al., 2008). Social experience also resets the circadian state and alters the activity of the clock neurons in *Drosophila* (Kim et al., 2013; Levine et al., 2002), which might drive the pleiotropic effects on diverse behavioral and physiological responses. Socially isolated flies sleep less than grouped flies, and this is mediated by contributions from dopaminergic modulation, cAMP signaling, and memory-related genes (Ganguly-Fitzgerald et al., 2006; Li et al., 2021). The influence of social experience extends to alterations in social behaviors such as courtship and aggression. *Drosophila melanogaster* males raised in groups display less intense courtship behavior toward female targets than isolated males (Dankert et al., 2009; Goncharova et al., 2016). In addition, *Drosophila melanogaster* males with prior exposure to mated females learn to reject mated females compared to males without such experience (Dukas, 2005). Social isolation has been shown to consistently increase aggression in various *Drosophila* species, both in males and females, including *D. melanogaster* (Agrawal et al., 2020; Liu et al., 2011; Ueda and Kidokoro, 2002; Ueda and Wu, 2009; Wang et al., 2008) and *D. suzukii* (Belenioti and Chaniotakis, 2020). The shared effects of social experience, combined with an unmatched set of genetic tools for labeling and manipulating neurons, make *Drosophila* an excellent model to explore molecular and circuit mechanisms by which social experience alters gene expression, circuit function, and behaviors.

Genes and circuits regulating *Drosophila* social behaviors, like courtship and aggression, have been well established. Two transcription factors, Fruitless (Fru) and Doublesex (Dsx), play crucial roles in governing the development and function of social circuits in *Drosophila*. Fru establishes most of the circuits mediating courtship and aggression (Anand et al., 2001; Billeter et al., 2006; Dickson, 2008; Koganezawa et al., 2016; Neville et al., 2014; Vrontou et al., 2006; Yamamoto and Koganezawa, 2013). The *fruitless* locus generates multiple alternative splice isoforms for RNAs transcribed from seven promoters (Billeter et al., 2006; Dalton et al., 2013; Demir and Dickson, 2005; Goodwin et al., 2000; Lee et al., 2000; Neville et al., 2014; von Philipsborn et al., 2014). The transcripts from *fru* P1 promoter are alternatively spliced between males and females by Transformer (Tra), where the male isoforms (*fru^M^*) encode functional proteins, whereas female isoforms (*fru^F^*) do not produce any protein product (Goodwin et al., 2000; Lee et al., 2000). Other *fru* promoters produce common isoforms in both male and female circuitry (Goodwin et al., 2000; Lee et al., 2000). Fru^M^ is expressed in an interconnected neural network throughout both the peripheral and central nervous system, and its expression is required for the development, function, and plasticity of the circuit that drives male-specific behaviors (Billeter et al., 2006; Dalton et al., 2013; Demir and Dickson, 2005; Goodwin et al., 2000; Lee et al., 2000; Neville et al., 2014; von Philipsborn et al., 2014). Owing to the function of Fru^M^ in establishing the structural and functional dimorphism of social circuits, it is unsurprising that Fru^M^ is both necessary and sufficient for sex-specific behaviors like male courtship.

A second transcription factor, Dsx, is essential for sex specification and differentiation in various tissues (Burtis and Baker, 1989; Dauwalder, 2011; Hildreth, 1965; Rideout et al., 2010; Villella and Hall, 1996). Unlike *fru,* which exhibits a high degree of evolutionary divergence, *dsx* is molecularly and functionally conserved across invertebrates and vertebrates (Clough et al., 2014; Kopp, 2012; Murphy et al., 2007; Rideout et al., 2010). Dsx belongs to the DMRT (Doublesex and Mab-3 Related Transcription factor) family, which is involved in regulating sex-specific differentiation (Kopp, 2012; Murphy et al., 2007).

*Doublesex* is also differentially spliced in males (*dsx^M^*) and females (*dsx^F^*) in *Drosophila*, and shows sexually dimorphic expression in the central brain (Burtis and Baker, 1989; Dauwalder, 2011; Hildreth, 1965; Rideout et al., 2010; Villella and Hall, 1996; Waterbury et al., 1999), albeit in fewer neurons compared to *fru^M^*. Disrupting Dsx function or blocking the activity of *dsx*-positive neurons alters courtship in both males and females (Han et al., 2022; Pan and Baker, 2014; Rideout et al., 2010; Zhou et al., 2014). Dsx^M^ also mediates social experience-dependent courtship learning exhibited by *fru^M^* mutant males, suggesting Fru^M^ drives innate courtship behaviors, while Dsx^M^ has a more modulatory effect that can be influenced by social experience in the absence of Fru^M^ (Pan and Baker, 2014). This is likely driven by the redundant function of Dsx^M^ and Fru^M^ both of which are expressed in a set of central courtship command neurons (P1 neurons) in males, which are causally linked to initiation of courtship (Inagaki et al., 2014; Kimura et al., 2008; Pan et al., 2012; Von Philipsborn et al., 2011).

Previous studies have shown that social context reprograms the expression and chromatin state of *fru* and *dsx* in pheromone-sensing neurons and alters the responses of these neurons to modify behaviors in a *fru^M^*-dependent manner (Deanhardt et al., 2023; Hueston et al., 2016; Sethi et al., 2019; Zhao et al., 2020). These findings highlight an intimate connection between social experience, chromatin remodeling, transcriptional modulation, and neuronal activity in the peripheral nervous system. Yet, it remains unclear how gene expression and neuronal activity are regulated in the central brain in response to social experience, and how these changes contribute to behavioral plasticity, particularly in male courtship.

Here, we investigated the impact of social experience on gene expression and neuronal activity in the central nervous system and how it reprograms male courtship behaviors in *Drosophila melanogaster.* We showed that group housing suppresses both evoked neural responses in courtship circuits and male courtship vigor, in agreement with previous research causally linking the activity of P1 neurons and courtship behaviors (Inagaki et al., 2014; Kimura et al., 2008; Pan et al., 2012). To identify differentially expressed genes responding to social experience, we carried out bulk RNAseq from the whole brain and single-cell RNAseq from *fru^M^*– and *dsx^M^*-positive neurons in the brains of grouped and isolated males, which revealed three major findings: 1) Social isolation increases *fru^M^* and *dsx^M^*reporter expression, the expression of *fru* in clock neurons, and *dsx* throughout the brain. Manipulating Fru^M^ function in different neuronal populations within social circuits has differential effects on social experience dependent modulation of courtship. 2) Group housing increases the expression of immediate early genes like *sr* and *Hr38* throughout the neurons within social circuits. The knockdown of *sr* in *fru*-positive neurons derepresses courtship in group-housed males to levels seen in isolated males. 3) Social experience alters the expression of Fru^M^/Dsx^M^ target genes encoding circadian regulators (*i.e. Clk*, *cry*) and regulators of male mating rhythm (*Sik3*) in the brain, which causally drive courtship responses in group-housed males. Overall, our results suggest that social experience reprograms Fru^M^ transcriptional cascades and their downstream target circadian genes within the social circuits to modulate courtship vigor.

## Results

### Social experience induces plastic changes in male fruit fly courtship behaviors

To assess how social experience influences male courtship behaviors, we examined the courtship activity of wild-type *Canton-S* (*CS*) male flies that were either group-housed (GH) or single-housed (SH) for seven days starting right after eclosion (Fig. 1). Males kept in groups exhibited significantly lower courtship intensity toward virgin females and exhibited suppressed P1 courtship command neuron activity compared to those raised in isolation (Fig. 1A-D; Supplementary Fig. 1B-D). Notably, the reduced courtship vigor in GH males was primarily due to an increased latency of the courtship onset (Fig. 1D). Once aroused, both GH and SH males courted females actively and persistently (Fig. 1A-D; Supplementary Fig. 1A).

**Figure 1.**
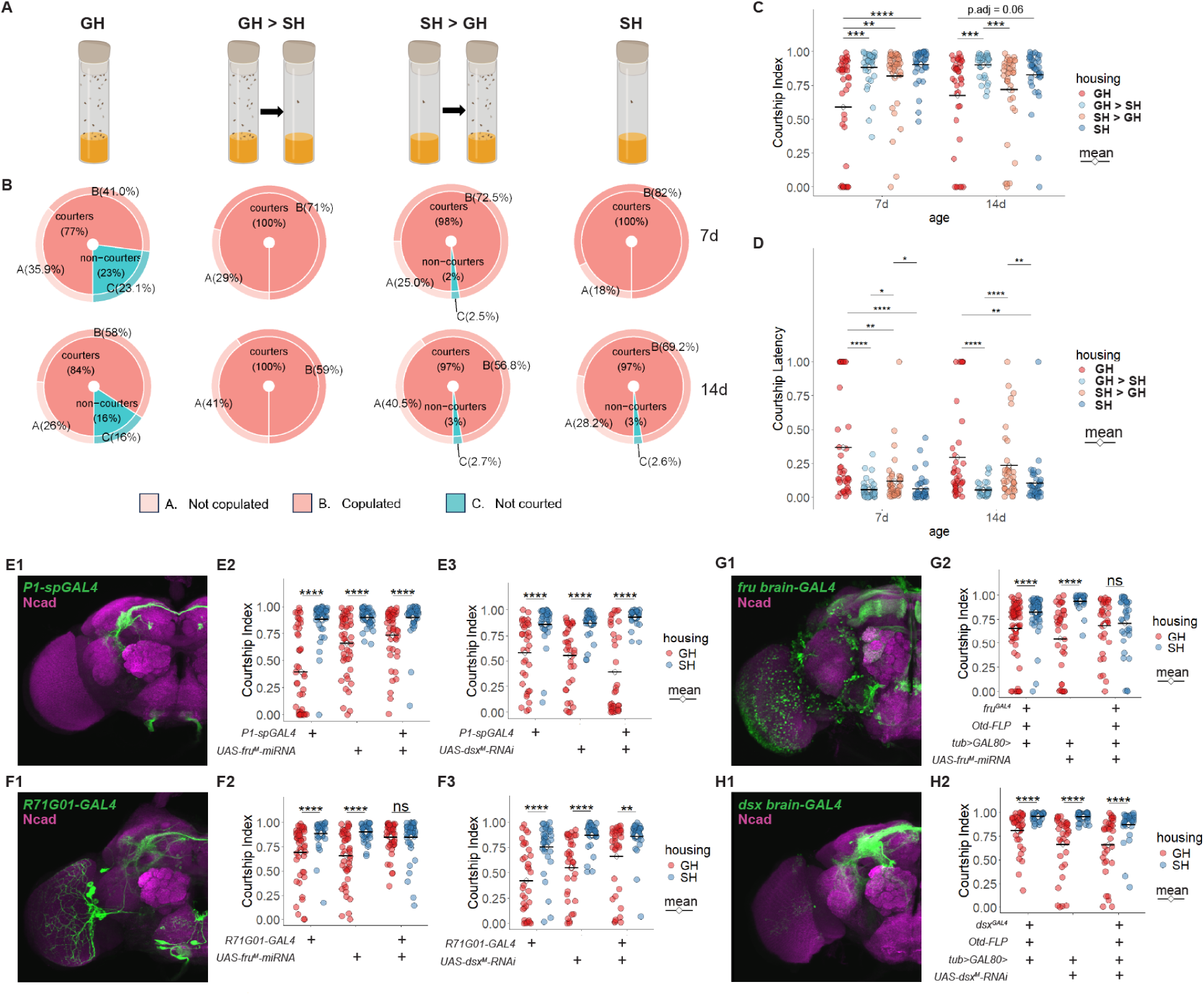
Social experience induces plastic changes in male fruit fly courtship behaviors and P1 neuronal activity. **(A)** Experimental paradigm for the setup of four different housing conditions. **(B)** Pie-Donut charts showing the percentage of flies at each courtship stage. **(C-D)** Courtship Index **(C)** and Courtship Latency **(D)** in male-female single pair assays with male flies raised under four different housing conditions. The Kruskal–Wallis rank sum test, followed by Dunn’s multiple comparisons test, was applied for the comparison among different housing conditions grouped by age. Significance of adjusted P-value (P_adj): *P_adj < 0.05, **P_adj < 0.01, ***P_adj < 0.001, ****P_adj < 0.0001. **(E1)** P1 neurons labelled by *P1-spGAL4> mCD8::GFP* in the brain, stained for GFP (green) and Ncad (magenta). See Methods for genotypes. **(E2-E3)** Courtship Index from male–female single-pair assays in control and experimental groups with *fru* **(E2)** or *dsx* **(E3)** knockdown in P1 neurons labeled by the stricter P1 driver. **(F1)** Neurons labelled by *R71G01GAL4> mCD8::GFP* in the brain, stained for GFP (green) and Ncad (magenta). **(F2-F3)** Courtship Index from male–female single-pair assays in control and experimental groups with *fru* **(F2)** or *dsx* **(F3)** knockdown in P1 neurons labeled by the broader P1 driver. **(G1)** Neurons labelled by *fru brain-GAL4> mCD8::GFP* in the brain, stained for GFP (green) and Ncad (magenta). **(G2)** Courtship Index from male–female single-pair assays in control and experimental groups with *fru* knockdown in *fru* positive neurons in the brain labeled by the intersection of *fru* and *Otd*. **(H1)** Neurons labelled by *dsx brain-GAL4> mCD8::GFP* in the brain, stained for GFP (green) and Ncad (magenta). **(H2)** Courtship Index from male–female single-pair assays in control and experimental groups with *dsx* knockdown in *dsx* positive neurons in the brain labeled by the intersection of *dsx* and *Otd*. The Wilcox test was applied to the comparison between GH and SH conditions of the same genotype. Significance of adjusted P-value (P_adj): *P_adj < 0.05, **P_adj < 0.01, ***P_adj < 0.001, ****P_adj < 0.0001.

To determine whether these socially induced behavioral changes are reversible, we tested the courtship behaviors in four different conditions: continuous GH, GH followed by SH (GH > SH), SH followed by GH (SH > GH), and continuous SH, across 7– and 14-day durations, with the switch occurring at 3.5 days and 7 days, respectively (Fig. 1A-D; Supplementary Fig. 1A). Across both time frames, males housed entirely in groups consistently courted less vigorously than those housed entirely in isolation (Fig. 1A-D; Supplementary Fig. 1A). When males were transferred from GH to SH (GH > SH), their courtship behavior shifted to resemble that of SH males in both 7 and 14 days (Fig. 1A-D; Supplementary Fig. 1A), suggesting a restoration of courtship drive upon social isolation. Conversely, the effects were more complex when flies were first isolated and then grouped (SH > GH). We found that courtship is suppressed in males housed in groups compared to isolation for either 7 or 14 days. Switching from SH to GH halfway through the 7-day housing duration led to a slight decline in courtship activity (Fig. 1A-D; Supplementary Fig. 1A). Yet, switching to GH on day 7 led to courtship levels resembling GH levels by the end of 14 days (Fig. 1A-D; Supplementary Fig. 1A). This indicates that the effect of isolation is more dominant and that the suppressive effect of group housing accumulates over time. These results overall point to the reversible impact of social experience on courtship behavior, with isolation exerting a stronger enhancing effect and prolonged grouping exerting a more substantial suppressive effect.

To further examine the effect of population density on courtship behavior, we compared the courtship index of 7-day-old isolated males with that of males raised in groups of 5, 10, 15, 20, or 30 (Supplementary Fig. 1E). The results showed that even the smallest group size (5 males) was sufficient to suppress courtship behavior, and increasing group size beyond that did not produce significantly stronger suppression (Supplementary Fig. 1E). This suggests that social cues from a relatively small number of conspecifics are enough to modulate male courtship behavior.

### Fru^M^, but not Dsx^M^, mediates social experience-dependent changes in courtship vigor

Fru^M^ and Dsx^M^ are transcription factors in the sex-determination pathway and regulate the development and function of circuits for male courtship behaviors (Dauwalder, 2011; Rideout et al., 2010; Ryner et al., 1996). In the central brain, *fru^M^* is expressed throughout interconnected neurons, including the P1 courtship command neurons that co-express both *fru^M^* and *dsx^M^* (Kimura et al., 2008; Lee et al., 2000; Pan et al., 2012). Given the important function of Fru^M^ and Dsx^M^ in regulating courtship, we first investigated whether *fru^M^*or *dsx^M^* is required in P1 neurons for the social experience-dependent modulation of courtship. We performed RNAi knockdown of *fru^M^*or *dsx^M^* in P1 neurons using an intersectional stringent driver (*P1-spGAL4*, Fig. 1E, see Methods for genotypes) and a broader P1 driver (*R71G01-GAL4*, Fig. 1F). While *fru^M^*knockdown with the stringent driver did not alter courtship responses to social experience, knockdown with the broader P1 driver increased courtship in GH males to SH levels (Fig. 1E and F). Furthermore, knocking down *fru^M^* in *fru*-positive neurons in the central brain using an intersectional strategy (*fru* brain-GAL4, see Methods for genotypes) also eliminated the courtship difference, this time due to reduced courtship in SH males (Fig. 1G; Supplementary Fig. 2A). This is consistent with previous studies demonstrating that *fru^M^* is essential for male courtship behavior (Anand et al., 2001; Demir and Dickson, 2005; Pan and Baker, 2014; Ryner et al., 1996). Both *fru^M^* mutants and *fru^M^* knockdown in all *fru*+ cells result in significantly reduced courtship behaviors in both GH and SH conditions (Supplementary Fig. 2C and D). These findings suggest that, besides its canonical roles in the specification of male courtship behaviors, *fru^M^* might have diverse roles in different neuronal populations in reprogramming courtship behaviors in response to social experience.

RNAi-based knock down of *dsx^M^* in P1 neurons—using either the stringent or the broad P1 drivers (Fig. 1E and F)—and *dsx*-positive brain neurons (Fig. 1H; Supplementary Fig. 2B) did not alter the effect of social experience on courtship behavior. Although *dsx^0683/1649^*mutant males consistently courted less than wild-type controls in agreement with previous reports (Rideout et al., 2010; Villella and Hall, 1996), GH *dsx^0683/1649^*mutants still courted females less than their SH counterparts (Supplementary Fig. 2E). Thus, while *dsx^M^* affects normal male courtship and is necessary for keeping the high level of courtship, it might not mediate the modulation of courtship behaviors with social experience. Alternatively, the function of Dsx in mediating responses to social experience might reside in cells outside the brain, like head fat body cells (Clough et al., 2014). These results suggest that the majority of the effects of social experience on courtship are largely mediated by Fru^M^ but not Dsx^M^.

### Bulk and cell-type-specific single-cell RNA sequencing revealed transcriptional responses to social experience

Social experience-dependent changes in circuit function and courtship behaviors may be driven by alterations in gene expression in response to neural activity/inactivity during chronic social enrichment or isolation. Previous reports showed that, in the peripheral olfactory system, social experience and pheromone signaling alter *fru^M^* and *dsx^M^* expression, and the expression of many Fru^M^ target genes, including modulators of membrane potential (Deanhardt et al., 2023; Hueston et al., 2016; Sethi et al., 2019; Zhao et al., 2020). These transcriptional responses to social experience are accompanied by Fru^M^-dependent alterations in the pheromone responses of olfactory neurons (Sethi et al., 2019).

To identify differentially expressed genes in response to social experience, we performed bulk RNA sequencing on brain tissues from 7-day-old wild-type *Canton-S* (*CS*) males raised in grouped or isolated conditions (Fig. 2). Principal component analysis (PCA) revealed a clear separation between the two housing conditions across three biological replicates (Fig. 2A), indicating consistent transcriptomic differences. The expression of housekeeping genes remained stable between the two groups (Supplementary Fig. 3). Approximately 240 genes were differentially expressed (Fig. 2B, C), with roughly half upregulated in grouped males and the other half downregulated. Based on Gene Ontology terms (GO terms), these differentially expressed genes (DEGs) are involved in biosynthesis, metabolism, circadian rhythm, behavior, and defense responses (Fig. 2E-G). Notably, *dsx* showed a near-significant trend for reduction in transcript levels in GH males (Fig. 2D). In contrast, *fru* transcript levels did not show a significant difference between GH and SH males (Fig. 2D).

**Figure 2.**
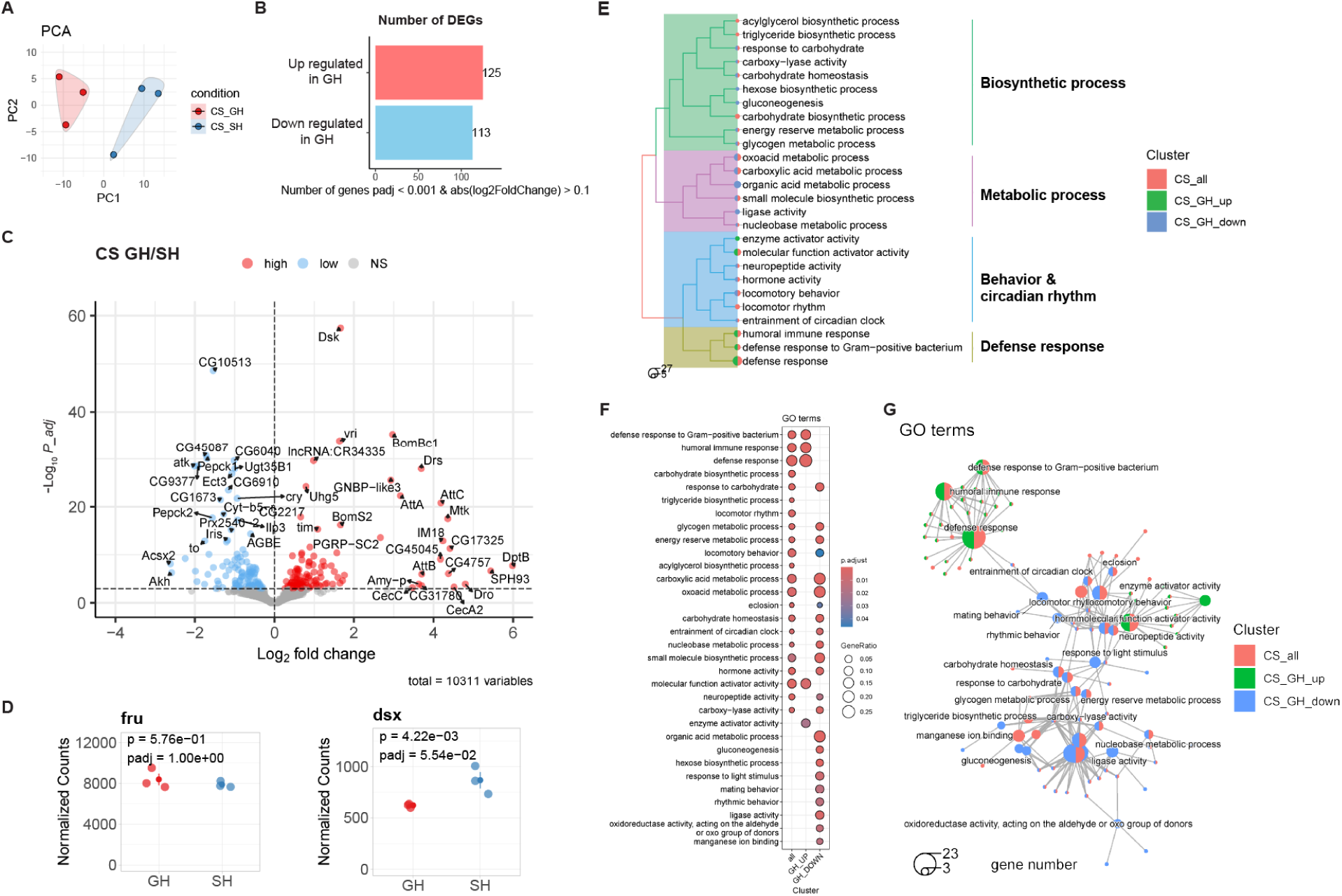
Bulk tissue RNA sequencing revealed diverse transcriptional changes in male brains to social experience. **(A)** PCA analysis of transcriptional profiles among biological replicates from brains of group-housed and single-housed wild-type *CS* male flies. **(B)** DEG numbers of GH/SH comparisons (adjusted P-value < 0.001, |log₂FC| > 0.1). Red indicates genes upregulated in the GH condition, and blue indicates genes downregulated in the GH condition. **(C)** Volcano plots displaying Log2FoldChange (GH/SH) and –Log10(P-adj) of genes. Red indicates genes upregulated in the GH condition, and blue indicates genes downregulated in the GH condition. **(D)** Transcript levels for *fru* and *dsx* among all male brain samples. **(E)** Summary of GO terms enriched in DEGs from the GH CS vs. SH CS comparison in male brains. Colors of branches indicate categories of GO terms. Colors of the pie indicate clusters of DEGs. The size of the pie indicates gene numbers. **(F)** Top 20 most significantly enriched GO terms for DEGs of GH/SH comparisons (q-value < 0.05). **(G)** Connection plots of the top 20 most significantly enriched GO terms for DEGs from GH/SH comparisons (q-value < 0.05). Big pies are GO terms, and small dots indicate different genes. Colors indicate clusters of DEGs. The size of the pie indicates gene numbers.

While bulk brain RNA sequencing provided insights into overall gene expression changes, it averages across all cells, masking small but meaningful changes within specific cell populations. To explore how social experience modulates gene expression at the cellular level, we performed single-cell RNA sequencing (scRNA-seq) on a combined pool of *fru^M^*+ brain cells and *dsx^M^*+ brain cells sorted from 7-day-old males raised under either GH or SH conditions (Fig. 3A). Using GFP reporters and fluorescence-activated cell sorting (FACS), we selectively sorted GFP-labeled *fru^M^*+ cells and *dsx^M^*+ cells from male brains (Fig. 3A). Notably, the percentages of *fru^M^::GFP* and *dsx^M^::GFP* positive cells in FACS were consistently higher in SH males compared to GH males (Fig. 3B, Supplementary Fig.4A). When analyzed separately, both *fru^M^*+ and *dsx^M^*+ populations showed an increased GFP+ cell ratio in SH male brains (Fig. 3B, Supplementary Fig.4A).

**Figure 3.**
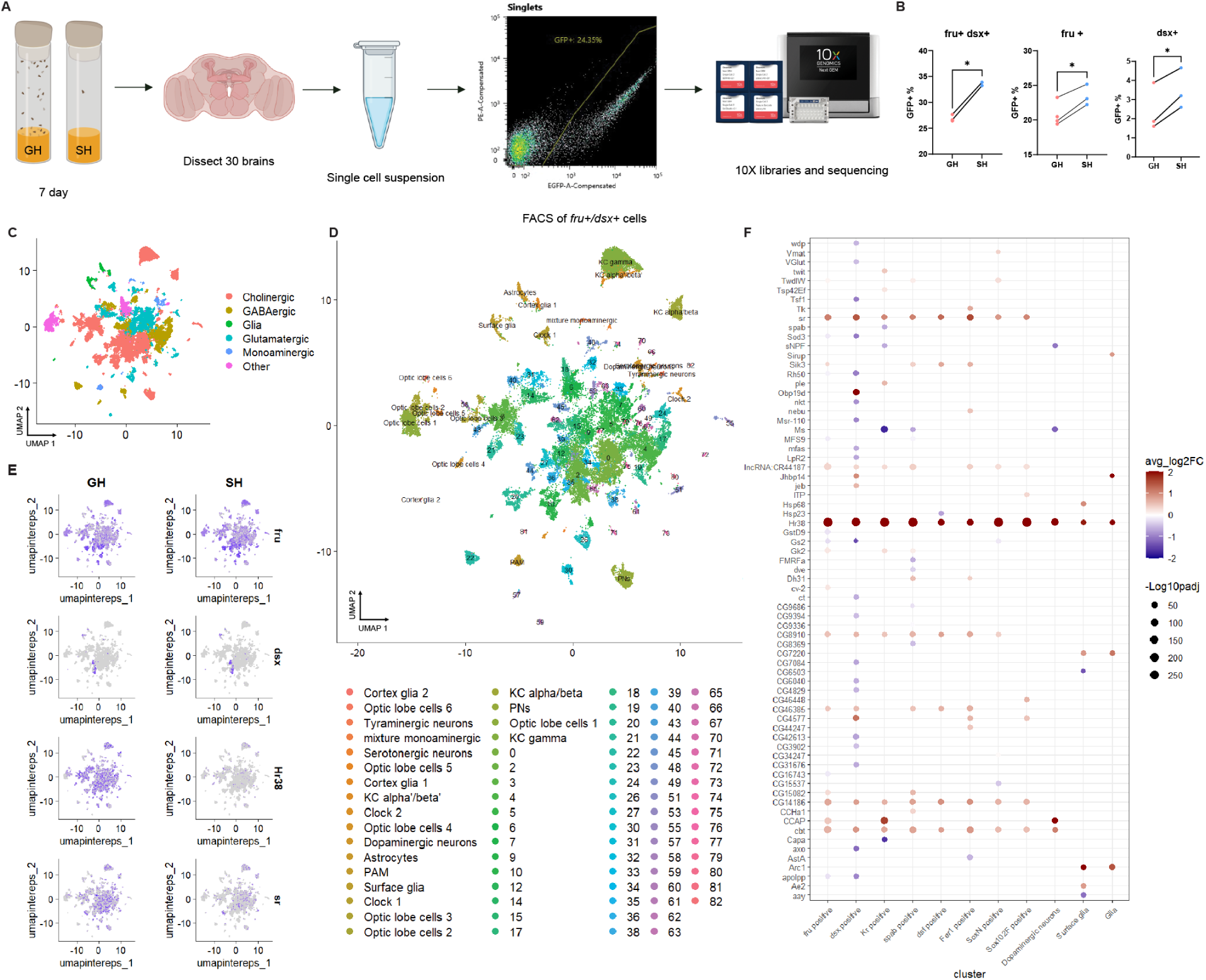
Cell-type-specific single-cell RNA sequencing revealed diverse transcriptional changes in male brains to social experience. **(A)** Workflow of the cell type-specific single-cell RNA sequencing of combined *fru*+ and *dsx*+ cells *(fru^GAL4^ 40XUAS-mCD8GFP; dsx-LexA 26XLexAop-mCD8GFP)* from brains of GH and SH males with the 10X Genomics platform. FACS: fluorescence-activated cell sorting. Created with BioRender. **(B)** Proportion of GFP-labeled combined *fru*⁺ cells and *dsx*^+^ cells, *fru*^+^ only cells, and *dsx*^+^ only cells in FACS. **(C-D)** Cluster annotation of cell-type-specific single-cell RNA sequencing. **(C)** UMAP plot illustrating six major cell classes, distinguishing glial cells from neurons characterized by their neurotransmitter profiles. **(D)** UMAP plot illustrating 83 specific cell clusters integrated across all brain samples. The annotations (bottom) were determined using marker gene expression. **(E)** Visualization of expression pattern of *fru*, *dsx*, *Hr38,* and *sr* in GH and SH male brains. Each dot is one cell colored by the expression levels of genes. **(F)** Pseudobulk analysis results showing Log2 fold changes (GH/SH) and –Log10 adjusted P-values of DEGs across cell subsets defined by specific marker genes, as indicated on the x-axis.

We then sought to determine whether this increase was restricted to particular cell types or distributed across many. At resolution 1, cells were grouped into 83 small clusters from six major clusters (Fig. 3C and D). All clusters expressed *fru^M^*, and some subsets expressed *dsx^M^*(Fig. 3E, Supplementary Fig.6A and B). Based on previously known gene markers (Davie et al., 2018; Park et al., 2022), we manually annotated 21 clusters (Fig. 3D). Overall, the proportions of these annotated cell types were similar between GH and SH samples, with clusters 6 and 18 showing the most significant variability across replicates, likely due to low RNA counts (Supplementary Fig. 4). Thus, they were excluded from further comparative analysis (Supplementary Fig. 5). Side-by-side comparisons of replicates 1 and 2 showed mostly consistent or near clustering proportions (Supplementary Fig. 5A and C). However, comparing proportions alone can obscure actual differences in cell abundance, as each sample’s proportions are normalized to add up to 1. To capture changes in cell numbers more accurately, we adjusted for the percentages of GFP+ cells from FACS data and applied these scaling factors to the cluster counts (Supplementary Fig. 5B and D). With this adjustment, many clusters from SH brains exhibited increased counts compared to GH brains (Supplementary Fig. 5B and D). This indicates that the rise in *fru^M^+/dsx^M^+* cells in SH male brains results from small increases, likely in transcriptional reporter activity, across many clusters, rather than expansion of a specific *fru^M^+* or *dsx^M^+* cell population.

Next, we conducted pseudobulk differential expression analysis across clusters to identify genes altered by social experience. Two immediate early genes (IEGs), *Hr38* (*Hormone receptor-like in 38*) and *sr* (*stripe*), known to respond to neural activity (Chen et al., 2016; Fujita et al., 2013; Takayanagi-Kiya et al., 2023), showed robust upregulation in most clusters (excluding glia) in GH brains compared to SH brains (Fig. 3E, Supplementary Fig. 6D and E). A third activity-responding gene, *cbt*, exhibited differential expression in only a few clusters (Supplementary Fig. 6E). These trends were largely consistent with the bulk RNA-seq data, where *Hr38* and *sr* also showed higher expression in GH brains, though *Hr38* did not reach statistical significance (Supplementary Fig. 6C). Given that *Hr38*, *sr,* and *cbt* showed strong upregulation in GH brains across multiple cell clusters, we asked if this contributes to courtship suppression using RNAi-mediated knockdowns in *fru*+ or *dsx*+ neurons under GH conditions. We found that *fru^GAL4^* or *dsx^GAL4^* driven expression of most RNAi transgenes against *Hr38* and *sr* in the brain increased courtship compared to controls in the first round of screening (Supplementary Fig. 9A and B). We next tested whether this increase is sufficient to eliminate the courtship difference between grouped and isolated males by analyzing *Hr38* and *sr* RNAi knockdown phenotypes in both GH and SH conditions with additional controls (Fig. 4). Knocking down *sr,* but not *Hr38,* in *fru^M^*+ neurons in the central brain (*fru* brain-GAL4) elevated courtship in grouped males, eliminating the courtship difference between GH and SH males (Fig. 4A). In contrast, knocking down *Hr38* or *sr* in *dsx^M^*+ neurons in the brain (*dsx* brain-GAL4) did not eliminate the effect of social experience on courtship difference between GH and SH conditions (Fig. 4B). In addition to the role of *fru^M^* in social modulation of courtship, these results also implicate transcriptional changes in immediate early genes like *sr* in *fru^M^* neuronal circuits in mediating courtship suppression in response to group housing.

**Figure 4.**
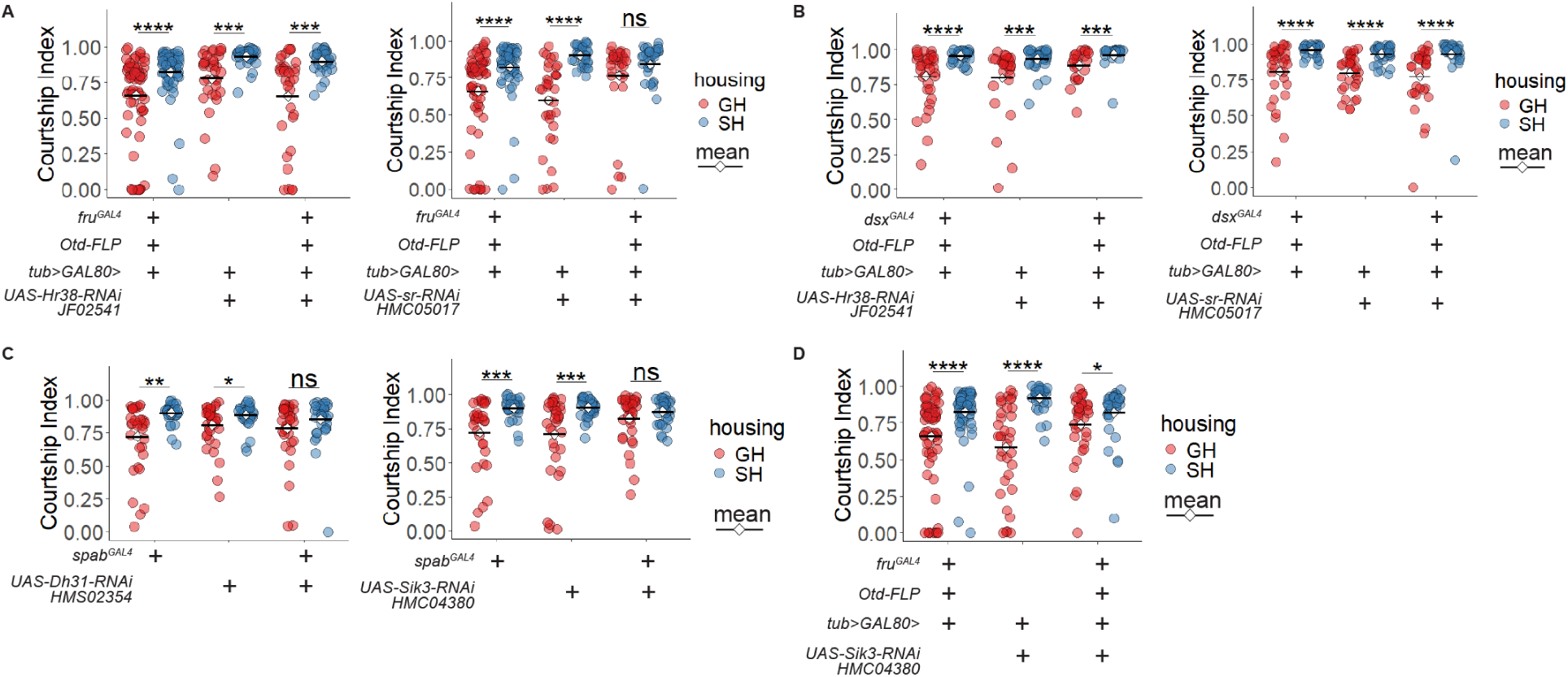
Knocking down *sr* or *Sik3* eliminated the courtship difference between GH and SH males. (**A-B**) Courtship Index from male–female single-pair assays in control and experimental groups with *Hr38* or *sr* knockdown in *fru* brain neurons **(A)** or *dsx* brain neurons **(B)**. **(C)** Courtship Index from male– female single-pair assays in control and experimental groups with *Dh31* or *Sik3* knockdown in *spab*+ cells. **(D)** Courtship Index from male–female single-pair assays in control and experimental groups with *Sik3* knockdown in *fru* brain neurons. The Wilcox test was applied to the comparison between GH and SH conditions of the same genotype. Significance of adjusted P-value (P_adj): *P_adj < 0.05, **P_adj < 0.01, ***P_adj < 0.001, ****P_adj < 0.0001.

As shown previously, P1 command neurons exhibit reduced evoked activity in GH males (Supplementary Fig. 1B-D). To determine transcriptional changes in P1 neurons that might underlie this reduction in activity, we also examined the DEGs in P1 neurons (Supplementary Fig. 10K). *Hr38* was again upregulated in GH P1 neurons. However, knocking down *Hr38* specifically in P1 neurons using *P1-spGAL4* did not affect the courtship response to group housing (Supplementary Fig. 10L), suggesting that P1 activity modulation could be driven by upstream circuit inputs rather than intrinsic transcriptional changes. Taken together, these results suggest that upregulation of immediate early genes within social circuits is associated with mediating the effects of group housing on courtship suppression.

### RNAi screening identifies candidate genes mediating social experience-dependent courtship plasticity

Single-cell RNAseq also revealed additional DEGs in various clusters, which are listed in Supplementary Figure 7. These DEGs are enriched in diverse biological processes, including neuronal signaling, neuromodulation, circadian regulation, and behavior (Supplementary Fig. 7B). To determine whether the DEGs identified in single-cell RNA-seq are causally involved in mediating social experience-dependent changes in courtship behavior, we conducted an RNAi screen. Even though some DEGs were in previously annotated clusters, like dopaminergic neurons or glia cells, there were still many DEGs in unannotated cells (Fig. 3D). Therefore, we used the FindMarkers function to identify marker genes for the four biggest clusters—excluding Cluster 1, which corresponded to mushroom body γ-lobe Kenyon cells with few DEGs (Fig. 3, Supplementary Fig. 8A).

Based on available GAL4 lines, we selected *Kr* and *spab* from Cluster 0, *dsf* and *Fer1* from Cluster 2, and *Sox102F* and *SoxN* from Cluster 3 as marker genes (Supplementary Fig. 8A). While none of these genes are exclusive to a single cluster, they served as valid entry points. We then performed pseudobulk DEG analysis in subsets expressing these markers, as well as in the *fru*+ and *dsx*+ subsets (Fig. 3F, Supplementary Fig. 8B). DEGs in these marker-defined populations were enriched in metabolism, signaling, and circadian processes, as well as behavior modulation (Supplementary Fig. 8C).

We used the cluster-specific marker GAL4 lines to knock down many of the DEGs and tested their effect on social modulation of courtship (Supplementary Fig. 9 and 10). Many DEGs within the *dsx*+ subset were from *dsx*+ glial cells (Supplementary Fig. 9C), whereas *dsx* brain-GAL4 does not target glia. These genes were thus tested genetically using *repo-GAL4* driven RNAi in glial cells (Supplementary Fig. 9 and 10). While many other RNAi lines showed no effect, a few genes, such as *Sik3* and *Dh31*, emerged as potent modulators of courtship plasticity, consistently showing elevated courtship in group-housed males when knocked down with multiple drivers and RNAi lines (Supplementary Figs. 9 and 10). Both genes are involved in circadian modulation of behaviors and are upregulated in specific cell clusters in the brain in response to group housing (Fig. 3F, Supplementary Fig. 7). *Sik3* knockdown in central brain *fru*+ neurons increased courtship in GH males and dramatically decreased the behavioral difference between GH and SH conditions (Fig. 4D). Similarly, knocking down *Sik3* in *spab*+ cells also diminished the GH– SH difference in courtship vigor by increasing courtship in grouped males (Fig. 4C). Meanwhile, even though knocking down *Dh31* in *spab+* cells eliminated the difference between GH and SH, the confidence was decreased due to the effects seen in controls (Fig. 4C). These findings implicate the circadian gene *Sik3* as an essential regulator of social experience-dependent modulation of courtship behaviors.

### Circadian genes mediate the effects of group housing on courtship behaviors

Previous studies have implicated *Sik3* in regulating the circadian male sex drive rhythm (Fujii et al., 2017), suggesting that social experience may influence internal arousal states by reprogramming the circadian system with downstream effects on the brain and behaviors. Thus, we revisited the DEGs in the bulk tissue RNAseq, where the GO terms centered on circadian rhythm-related processes (Fig. 2).

In the bulk brain RNAseq, three key circadian genes—*cryptochrome* (*cry*), *timeless* (*tim*), and *Clock* (*Clk*)—exhibited significant differential expression between GH and SH males (Fig. 2C). Interestingly, these four circadian genes (*Sik3, cry, tim* and *Clk*) are indicated as potential target genes transcriptionally regulated by Fru^M^/Dsx^M^ (Clough et al., 2014; Neville et al., 2014). The brain expression patterns of circadian genes *cry*, *Clk*, and *tim* in the clock network overlap, but are not always coexpressed (Fig. 5A and D). To confirm social experience-dependent transcriptional changes in *Clk, cry,* and *tim,* we analyzed their expression *in vivo* using existing transcriptional reporters and protein-GFP fusions in the brain (Fig. 5A and B, also see (Du et al., 2025)). Among the three circadian genes, we found that *Clk* transcript levels were upregulated in response to single housing (Fig. 5A and B). In contrast, we found Cry protein levels exhibited an increase in response to group housing (Fig. 5A and B). *tim,* on the other hand, did not show an overall difference in transcript levels (Fig. 5A and B). Overall, these results suggest that some circadian genes are modulated by social experience.

**Figure 5.**
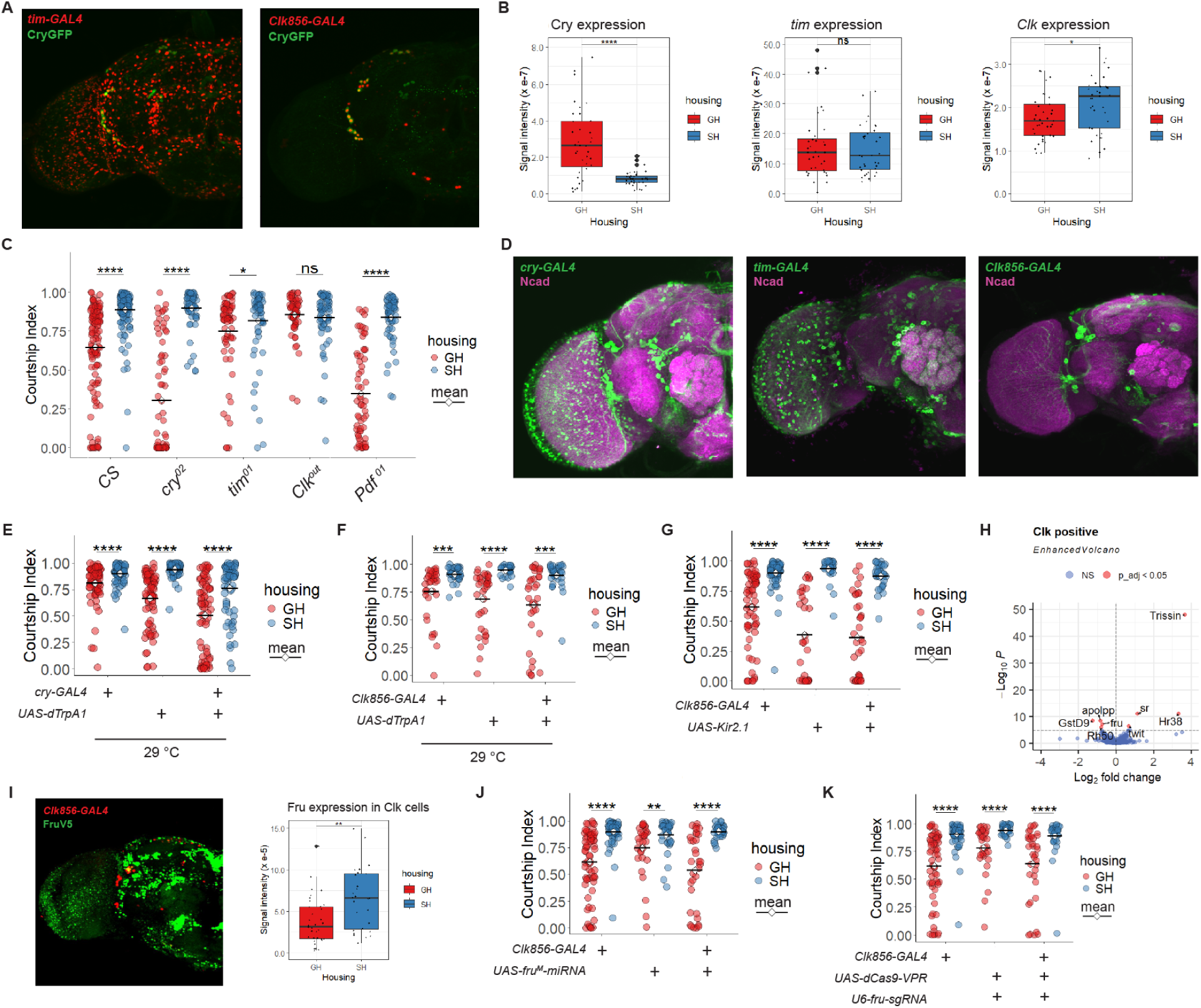
Circadian genes mediate the effects of group housing on courtship behaviors. **(A)** Cry+ cells labelled by GFP, with *tim*+ or *Clk*+ cells labelled by mCherryNLS in male brains, stained for GFP (green). **(B)** Quantification of fluorescence intensity of Cry labelled by GFP, and *tim*+ or *Clk*+ labelled by mCherryNLS in **(A)**. The sum of net fluorescence signals (fluorescence signals – background signals) was compared between GH and SH male brains. Unpaired t-test was applied. **(C)** Courtship Index from male– female single-pair assays in *CS*, *cry* mutants, *tim* mutants, *Clk* mutants and *Pdf* mutants. **(D)** The GFP expression pattern driven by *cry-GAL4*, *tim-GAL4* and *Clk856-GAL4* used for activation/silencing of neuronal activity, stained for GFP (green) and Ncad (magenta). **(E-F)** Courtship Index from male–female single-pair assays with activation of *cry+* **(E)** or *Clk*+ **(F)** neurons using dTrpA1. **(G)** Courtship Index from male–female single-pair assays with silencing of *Clk*+ neurons using Kir2.1. **(H)** Volcano plots displaying Log2FoldChange (GH/SH) and –Log10(P-value) of genes in *Clk*+ cells from the single-cell RNAseq. Red indicates DEGs from pseudobulk analysis. **(I)** Left: Fru+ cells labelled by V5, with *Clk*+ cells labelled by mCherryNLS in male brains, stained for V5 (green). Right: Quantification of fluorescence intensity of Fru labelled by V5 in *Clk*+ cells. The sum of net fluorescence signals (fluorescence signals –; background signals) was compared between GH and SH male brains. Unpaired t-test was applied. **(J-K)** Courtship Index from male–female single-pair assays comparing control and experimental groups of *fru* knockdown **(J)** or *fru* overexpression **(K)** in *Clk*-expressing cells. For all courtship behaviors, the Wilcox test was applied to the comparison between GH and SH conditions of the same genotype. Significance of adjusted P-value (P_adj): *P_adj < 0.05, **P_adj < 0.01, ***P_adj < 0.001, ****P_adj < 0.0001.

**Figure 6.**
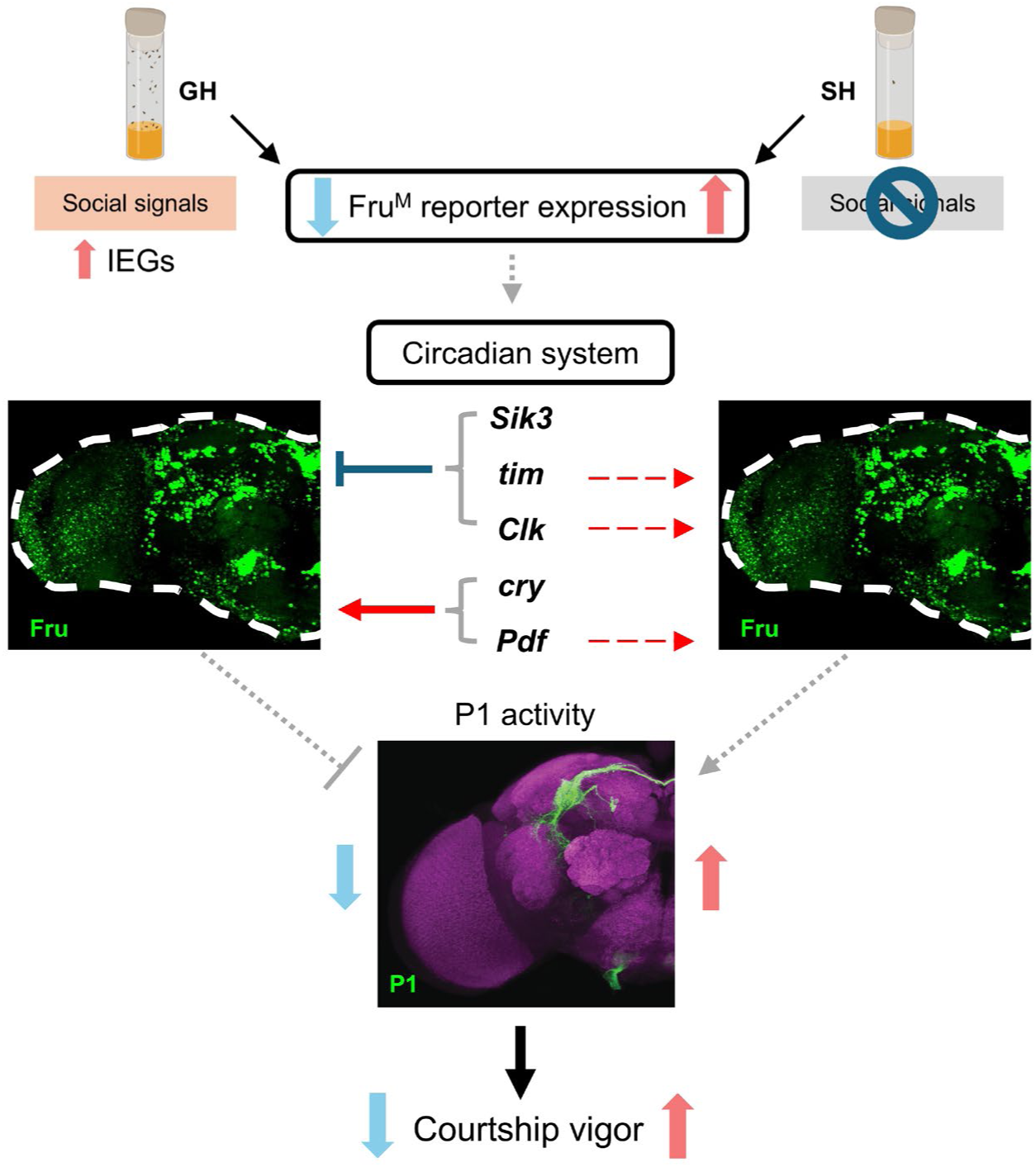
A model for social experience-dependent modulation of courtship. Group housing suppresses the courtship vigor of male flies. The transcription factor Fru^M^ plays an essential role in this modulation, acting within distinct neuronal subtypes to integrate social signals and modulate behavioral output. The increase in *fru^M^+* cells in the central circuits likely occurs as an increase in *fru^M^*expression, which drives an increase in courtship in response to social isolation. In group-housed conditions, upregulation of the immediate early gene, *sr,* in *fru*+ neural circuits mediates male courtship suppression. Circadian genes (*Sik3*, *tim, Clk, cry, Pdf*), potentially regulated by Fru^M^/Dsx^M^ transcription factors, also contribute significantly to social experience-dependent modulation in courtship behaviors. *cry* and *Pdf* function oppositely to *Sik3*, *tim* and *Clk* to modulate courtship behaviors in the GH condition. In contrast, the effects of these circadian genes are limited in socially isolated males, which might dominate the behavioral elevation by regulating additional factors. These changes all together lead to suppressed evoked P1 courtship command neuron responses and male courtship behaviors under GH condition relative to the SH condition.

To investigate whether changes in circadian gene expression with social experience contribute to experience-dependent changes in courtship behavior, we examined the courtship in *cry, tim,* and *Clk* mutants. We also investigated *Pdf* mutants, because of its coexpresssion and interaction with *cry* in a subset of clock neurons (Cusumano et al., 2009; Zhang et al., 2009), as well as its transcriptional downregulation in response to group housing (Supplementary Fig. 7A). Compared to wild-type *CS* males, *Clk* mutants completely eliminated the difference in courtship by increasing courtship in GH males (Fig. 5C; Supplementary Fig. 11B). *tim* mutants also showed a reduced difference in courtship between GH and SH conditions due to increased courtship in GH males (Fig. 5C; Supplementary Fig. 11B). In contrast, *cry* and *Pdf* mutants showed an accentuated courtship difference between GH and SH males, mostly due to further courtship suppression in GH males (Fig. 5C; Supplementary Fig. 11B). We also manipulated neuronal activity within clock neuron populations. Silencing or stimulating neural activity using *Clk-GAL4-*driven *UAS-Kir2.1* or *UAS-dTrpA1* expression did not alter social experience-dependent changes in courtship behaviors (Fig. 5D, F and G). *tim-GAL4* driven *UAS-Kir2.1* or *UAS-TrpA1* expression disrupted viability (Fig. 5D). While *cry-GAL4* driven *UAS-Kir2.1* did not yield viable flies, interestingly, activating neuronal function using *cry-GAL4* driven *UAS-dTrpA1* mimicked *cry* and *Pdf* mutants in courtship responses to grouping (Fig. 5C-E; Supplementary Fig. 11C).

As *cry*, *tim*, and *Clk* are partially coexpressed with *fru^M^* in subsets of neurons in the brain that label the clock neural network, we further explored how gene expression is altered in these overlapping populations. Given the strong behavioral phenotype of *Clk* mutants, we focused on the *Clk*+ cell population and analyzed DEGs within this subset using single-cell RNA-seq data (Fig. 5H, Supplementary Fig. 11A). We observed that in group-housed males, *Hr38* and *sr* were upregulated in *Clk*+ cells, while *fru* was significantly downregulated in this population (Fig. 5H). We validated this expression level by quantifying Fru^M^ (FruV5) in *Clk*-expressing cells *in vivo* using imaging (Fig. 5I). To test whether these changes contribute to courtship modulation, we manipulated the expression of *Hr38*, *sr*, and *fru^M^* in *Clk*+ cells, followed by courtship assays. Specifically, we knocked down *Hr38*, *sr* (Supplementary Fig. 11D), and *fru^M^*, and overexpressed *fru^M^* in *Clk*+ neurons (Fig. 5J-K). However, none of these manipulations affected the courtship difference between GH and SH males compared to controls. Together, these results suggest that while circadian genes like *Clk*, *tim, pdf, cry Sik3*, are crucial for mediating the effect of group housing on courtship suppression, *fru^M^* and IEG function in *Clk*-positive neurons and their neuronal activity are not implicated. Rather, Fru^M^ function in other clock neurons (i.e, *Pdf* or *cry*-expressing neurons included in broad P1 drivers in Fig.1) might modulate courtship suppression in group-housed males. Yet, given the *Clk* mutant phenotype, these results raise the possibility that non-neuronal *Clk-*positive cells around the brain, like fat body cells or glia cells (Patop et al., 2023), might still have an influence on social modulation of behaviors. Overall, our results suggest that group housing might impinge upon the circadian networks serving as a general hub for reprogramming the arousal state or responsiveness of the core courtship circuits in the brain.

## Discussion

Social experiences, from isolation to social enrichment, exert profound and enduring influences on cognition, physiology, and behavior (Cacioppo and Hawkley, 2009; Dow-Edwards et al., 2019; Jeste et al., 2020). Loneliness is associated with an increased incidence and severity of numerous neuropsychiatric and neurodegenerative disorders, highlighting its emergence as a contemporary epidemic, particularly detrimental to elderly populations who are more susceptible to social isolation epidemic (Cacioppo and Hawkley, 2009; Dow-Edwards et al., 2019; Jeste et al., 2020). Despite these widely known effects of social isolation on brains and behaviors, the molecular and circuit-based mechanisms through which social experiences shape health and behavior remain largely unexplored. In this study, we used *Drosophila melanogaster* to investigate the neural and molecular pathways by which social experiences influence male courtship behavior.

Group housing suppresses male courtship activity and reduces the evoked responses in P1 courtship command neurons. The transcription factor Fru^M^ plays an essential role in this modulation, acting within distinct neuronal subtypes to integrate social signals and modulate behavioral output. Furthermore, the immediate early gene *sr*, functioning in *fru*-expressing neural circuits, are critical for maintaining the behavioral distinctions between grouped and isolated males. Beyond *fru^M^*, circadian genes (*Sik3*, *tim, cry, Clk* and *Pdf*) – some are putative Fru^M^ targets (Neville et al., 2014) – also contribute significantly to sustaining the social context-dependent variation in courtship behavior. Given that social experience resets the circadian rhythm and alters the activity within central clock neurons (Kim et al., 2013; Levine et al., 2002), our results highlights modulation of circadian state as a causal mechanism through which social experience influences courtship behaviors.

Previous studies have shown that monosexual group housing suppresses male courtship behaviors, such as wing extension and courtship song (Dankert et al., 2009; Goncharova et al., 2016). Goncharova et al. further proposed that pre-grouped males avoid contact during courtship tests, possibly due to a conditioned fear response arising from male-male aggression during group housing. In the present study, we confirmed that group housing suppresses courtship vigor; however, this suppression is reversible— isolating previously grouped males restores their courtship activity. Notably, the degree of suppression increases with prolonged group housing, suggesting a time– and exposure-dependent mechanism underlying this behavioral plasticity.

At the neural circuit level, P1 neurons are key command neurons for male courtship behavior, capable of sustaining courtship-associated states for minutes (Inagaki et al., 2014; Jung et al., 2020; Kimura et al., 2008; Pan et al., 2012; Von Philipsborn et al., 2011). Previous studies found that isolated males were more sensitive to the same level of optogenetic stimulation of P1 neurons and showed an increase in wing extension compared to grouped males in response to the stimulation (Inagaki et al., 2014). *Ex vivo* imaging of P1 neurons in grouped male brains showed lower evoked responses than isolated male brains (Inagaki et al., 2014; Zhao et al., 2024). These results suggest that P1 neuronal activity less sensitized in GH male brains. Consistent with these findings, in this study, we show that P1 neurons in live-behaving grouped males exhibited reduced evoked responses to female cues. Together, these data support the idea that group housing diminishes P1 neuron sensitivity, thereby increasing courtship latency and reducing overall courtship vigor.

The question then arises: how does social experience regulate the activity of P1 neurons? As an integrative hub for multiple sensory inputs, P1 neurons are modulated by upstream circuit components (Clowney et al., 2015; Kohatsu et al., 2011; Nässel and Wu, 2022). Social conditions likely alter the activity or connectivity of these upstream inputs. For instance, Drosulfakinin (DSK) has been shown to inhibit male courtship through its receptor CCKLR-17D3 on P1 neurons, and group housing increases synaptic connections between DSK-expressing neurons and P1 neurons in male brains (Wu et al., 2019). This enhanced inhibition may underlie the reduced courtship vigor observed in grouped males. In the future, monitoring responses of individual neurons upstream of P1 in the courtship circuit in grouped or isolated male brains will reveal neuronal paths implicated in the modulation of behaviors with social experience.

Given that P1 neurons express both *fruitless^M^*and *doublesex^M^*, it is important to determine whether these courtship regulators contribute to the social experience-dependent modulation of courtship. Single-cell and bulk RNAseq experiments revealed that social isolation increases *fru^M^* and *dsx^M^* reporter expression, the levels of *fru^M^*transcripts in *Clk+* cells, and levels of *dsx^M^* transcripts throughout the brain. Even though bulk RNAseq experiments showed that *dsx^M^* expression increases in isolated male brains, it is not required in neurons for driving the courtship difference between grouped and isolated males. This suggests that *dsx^M^*may influence other behaviors or structural changes in the brain rather than courtship per se, or its function in this process is located in non-neuronal cells around the brain. In contrast, knocking down *fru^M^* in *fru^M^+* neurons in the brain abolished the behavioral differences between grouped and isolated males, by decreasing courtship in isolated males. This result is in agreement with a role for Fru^M^ in positively regulating courtship. Interestingly, knocking down *fru^M^* in certain neurons increases courtship in grouped males (Fig. 1F), suggesting Fru^M^ might play different roles in various neuronal populations with opposing influences on courtship circuits. Regardless, these results demonstrate that Fru^M^ function is essential for encoding the behavioral impact of social experience.

Beyond *fru^M^*, this study also identified circadian-related genes, such as *Clk*, *tim*, *Pdf, cry,* and *Sik3*, as key modulators of courtship behavior under different social contexts. *Sik3* expression, which we found to be upregulated in group-housed brains, has been shown to influence male sexual circadian rhythms (Fujii et al., 2017). Mutants for *tim* and *Clk* exhibited disrupted or eliminated differences in courtship between grouped and isolated males, indicating that circadian reprogramming likely contributes to social modulation of courtship. Compared to *Clk* and *tim* mutants, *cry* mutants further decrease courtship responses in grouped males, suggesting the effects of circadian genes on reprogramming behaviors are restricted to group-housed conditions. It is interesting that some of these circadian genes have binding sites for Fru^M^ or Dsx^M^ in their upstream regulatory elements (Clough et al., 2014; Neville et al., 2014), suggesting potential transcriptional cascades in response to social experience. Future experiments will reveal the cells where these regulatory interactions occur and how activity/inactivity within courtship circuits induced through social experience affects their expression.

Within the *Clk*+ cells, *sr* and *Hr38* expression levels were upregulated in grouped male brains compared to isolated brains, whereas *fru* expression was decreased. Yet, manipulating *fru^M^*, *sr*, or *Hr38* expression in *Clk*+ neurons did not impact courtship behavior, suggesting that they do not function in *Clk+* cells for grouping-induced suppression of courtship. A recent study showed that neurons where *Clk* and *fru^M^* overlap influence courtship, and that *fru^M^*+ neurons receive input from *Clk*+ neurons (Deluca et al., 2025). These findings suggest that while *Clk*+ neurons may not be part of the core courtship circuit, they can indirectly influence the function of neurons within courtship circuits. *Clk* mutants exhibited elevated courtship behavior in grouped males compared to wild-type, yet manipulating *Clk-*positive neural activity does not have an effect. One possible explanation is that in group-housed male brains, Clk protein functions in non-neuronal cells (i.e. fat body or glia) to suppress the function of the courtship circuits. In contrast, both manipulating the function of *cry* and *Pdf,* normally co-expressed in a subset of clock neurons, or the function of *cry-*positive neurons, resulted in further suppression of courtship in group housed males. These results suggest opposing function of different circadian genes and the activity of a subset of clock neurons mediate the effects of social experience on courtship responses to group housing.

Even though grouping-induced courtship suppression is disrupted in *Clk* and *tim* mutants, their mode of transcriptional modulation by social experience and their impact on courtship are not straightforward. Among the three core circadian genes, *Clk* expression consistently shows a downregulation in response to group housing using multiple approaches. In contrast, the expression of *tim* and *cry* shows some variation based on the methodology used. Given the complex transcriptional and translational regulation of the core circadian molecular oscillator, a deeper investigation into the social modulation of the circadian clock will help resolve these discrepancies.

One surprising finding we observed is that for both *Clk* and *cry* genes, the phenotypic trends tend to manifest in opposition to predicted outcomes based on transcriptional responses to social experience. For example, the *Clk* gene is upregulated in isolated male brains, while *Clk* mutants increase courtship in group-housed males. Similarly, Cry is upregulated in group-housed male brains, while *cry* mutants further decrease courtship in response to group housing. These are possibly due to the existence of double-negative regulatory loops regulating the function of clock neural networks by the circadian protein. Given that the phenotypic effect of circadian mutants is restricted to GH males, it is also possible that they contribute much less to the modulation of behavior in response to socially isolated conditions. These results support the hypothesis that circadian genes play essential roles in mediating the effects of social experience on male courtship behavior, agreeing with previous reports on reprogrammed circadian state with social experience (Levine et al., 2002). They also point to opposing effects of Cry and Pdf versus Tim and Clk in mediating the effects of group housing on courtship behaviors, and that there might be more dominant programs mediating behavioral response to social isolation that rely on Fru^M^ function.

Within both *Clk*+, *fru^M^*+, and *dsx^M^*+ neuronal subsets, the immediate early genes *sr* and *Hr38* showed elevated expression in grouped male courtship circuits. Functional experiments revealed that upregulation of *sr* within *fru^M^*+ neurons in response to group housing is required to suppress courtship. While it does not appear to act within P1 or *Clk*+ neurons directly, it may influence specific, yet unidentified, subsets of *fru*+ neurons or other clock neuron subsets. Future studies are needed to map the precise cellular substrates through which *sr* operate to mediate socially driven behavioral changes.

Beyond courtship, social experience modulates a wide array of behaviors in *Drosophila*, including sleep, feeding, locomotion, and aggression (Agrawal et al., 2020; Dankert et al., 2009; Ganguly-Fitzgerald et al., 2006; Goncharova et al., 2016; Kacsoh et al., 2018; Li et al., 2021; Ueda and Kidokoro, 2002; Wang et al., 2008; Zhao et al., 2024). The social isolation-induced rise in aggression is suggested to be regulated by genes involved in reactive oxygen species (ROS) metabolism, *Hyperkinetic* (*Hk*) and *glutathione S-transferase-S1* (*gsts1*) (Ueda and Wu, 2009), as well as the neuropeptide Drosulfakinin (Dsk) in the central brain (Agrawal et al., 2020), and cytochrome gene *Cyp6a20* in the peripheral olfactory systems (Wang et al., 2008). Furthermore, Or65a olfactory receptor neurons (ORNs), which respond to the male pheromone 11-*cis*-vaccenyl acetate (cis-VA), mediate the suppression of aggression in group housed male (Liu et al., 2011). Group-housed *Drosophila* males also decrease their locomotion compared to single-housed males, and this is mediated by P1 neurons and other *dsx^M^*-positive neurons, where neuropeptides for rhythmic locomotion and aggression, DH44 and TK, oppositely regulate locomotion through feedback loops (Zhao et al., 2024).

Whether a shared molecular or circuit-level mechanism underlies the effects of social experience across these diverse behaviors remains an open and compelling question. Given the pleiotropic suppressive effects of social experience on a variety of behaviors, there is potential for central arousal-related networks in mediating the effects. The finding that group housing induced courtship suppression requires circadian reprogramming is significant, as it might also underlie modulation of many of the behaviors in response to social experience, particularly in light of previous reports suggesting social experience resets circadian state (Levine et al., 2002). For example, many behaviors are suppressed during nighttime due to the modulation of arousal through molecular and circuit-level oscillations of the circadian clock (Duhart et al., 2023). Changes happening in the circadian system in response to social experience can provide a central modulatory hub that mediates many of the effects of social experience on organismal behaviors and physiology. It is likely that any change in circadian state would result in state changes in downstream neurons driving diverse behaviors. These can include P1 command neurons and dopaminergic neurons, as activity in both is affected by social experience and was shown to modulate courtship, aggression, feeding, and sleep (Agrawal et al., 2019; Alekseyenko et al., 2013; Ganguly-Fitzgerald et al., 2006; Karam et al., 2020; Zhang et al., 2018). A systematic examination of transcriptional and circuit mechanisms by which social signals engage arousal systems, like the circadian system, will be crucial for understanding the central strategies mediating the brain’s adaptive responses to social environments.

Social isolation has been increasingly recognized as a risk factor not only for mental health conditions such as depression and anxiety, but also for neurodegenerative diseases like Alzheimer’s (Cacioppo et al., 2015; Cacioppo et al., 2006; Cohen, 2004). By uncovering the mechanisms through which the social environment influences neural circuits, this study may help inform strategies to mitigate the negative consequences of social isolation on brain health across species, including humans. Thus, our findings contribute to a deeper understanding of how social experience shapes brain function and behavior.

## Methods

### *Drosophila* strains and husbandry

Flies were raised on standard fly food from Archon Scientific company at 25 °C (except the courtship assays with dTrpA1, labeled in figures) under a 12-h light: 12-h dark (LD) cycle. All experiments used male flies except targets of courtship behavior assays. Male flies used in courtship behavior assays, bulk tissue RNA sequencing, single-cell RNA sequencing, two-photon calcium live imaging and quantification of immunofluorescence and confocal microscope imaging were unmated and 7 days old unless mentioned otherwise. For male-female single-pair courtship behavior assays, unmated males were used as the test fly, and 4 to 6-day-old *w^1118^*virgin females were used as the target fly. Male flies used for courtship behavior assays were *w^+^* background (except *dsx* mutant) and did not have balancers. Fly stocks were obtained from the Bloomington Drosophila Stock Center (BDSC), the Vienna Drosophila Resource Center (VDRC), or kindly provided by other labs. Specific genotypes of males used in each figure are listed together with the sources for each fly stock in Table 1 (same sources are listed only once, at their first appearance). Split GAL4 lines *R15A01-AD* and *R71G01-DBD* were crossed together for intersectional labeling of P1 neurons (*P1-spGAL4*). *Otd-FLP/tub>GAL80>; fru^GAL4^* or *Otd-FLP; fru^GAL4^/tub>GAL80>* restricts expression from *fru^GAL4^*to neurons in the central brain (*fru brain-GAL4*) while *Otd-FLP/tub>GAL80>; dsx^GAL4^* restricts expression from *dsx^GAL4^* to neurons in the central brain (*dsx brain-GAL4*).

**Table 1.**
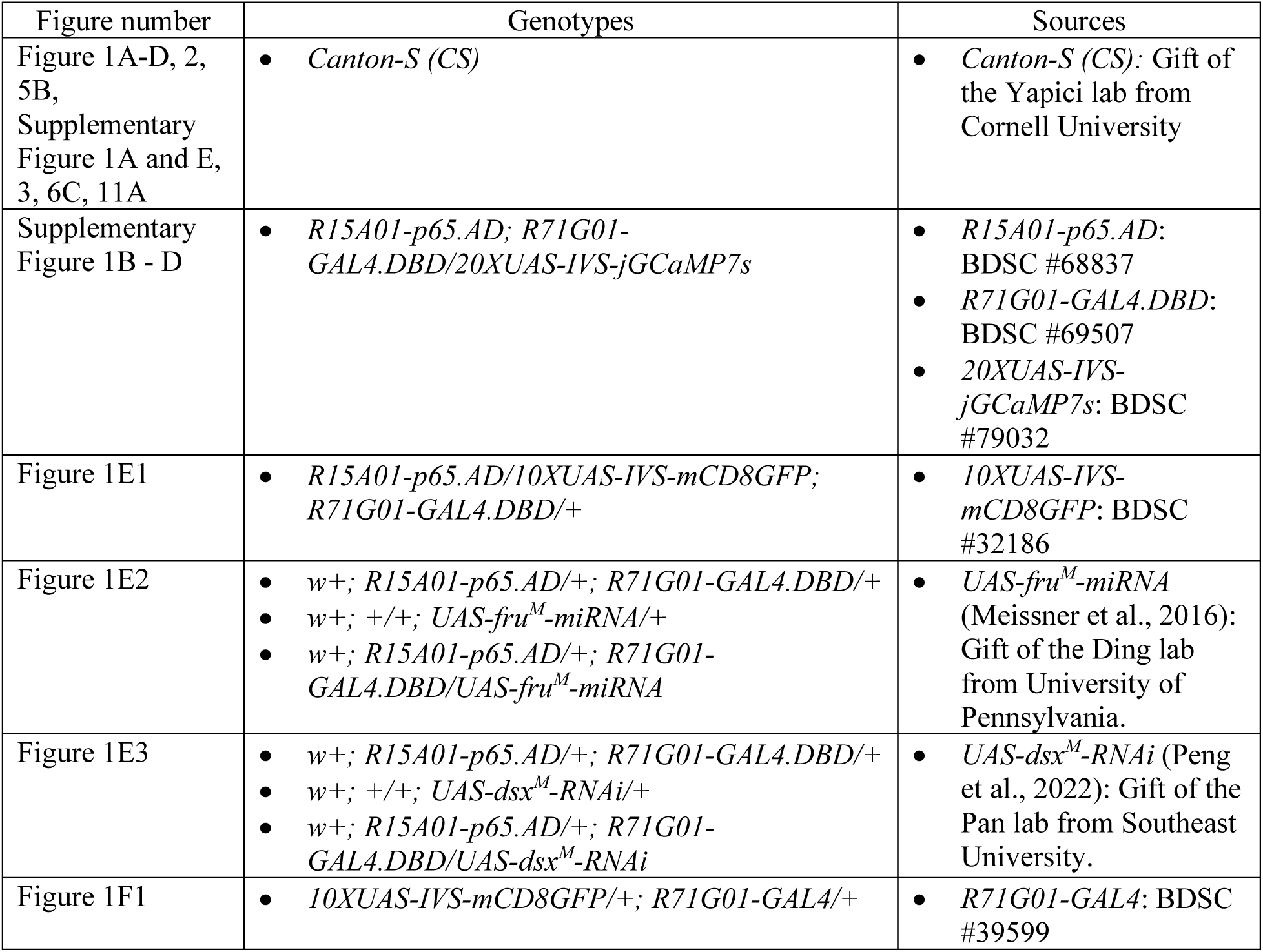
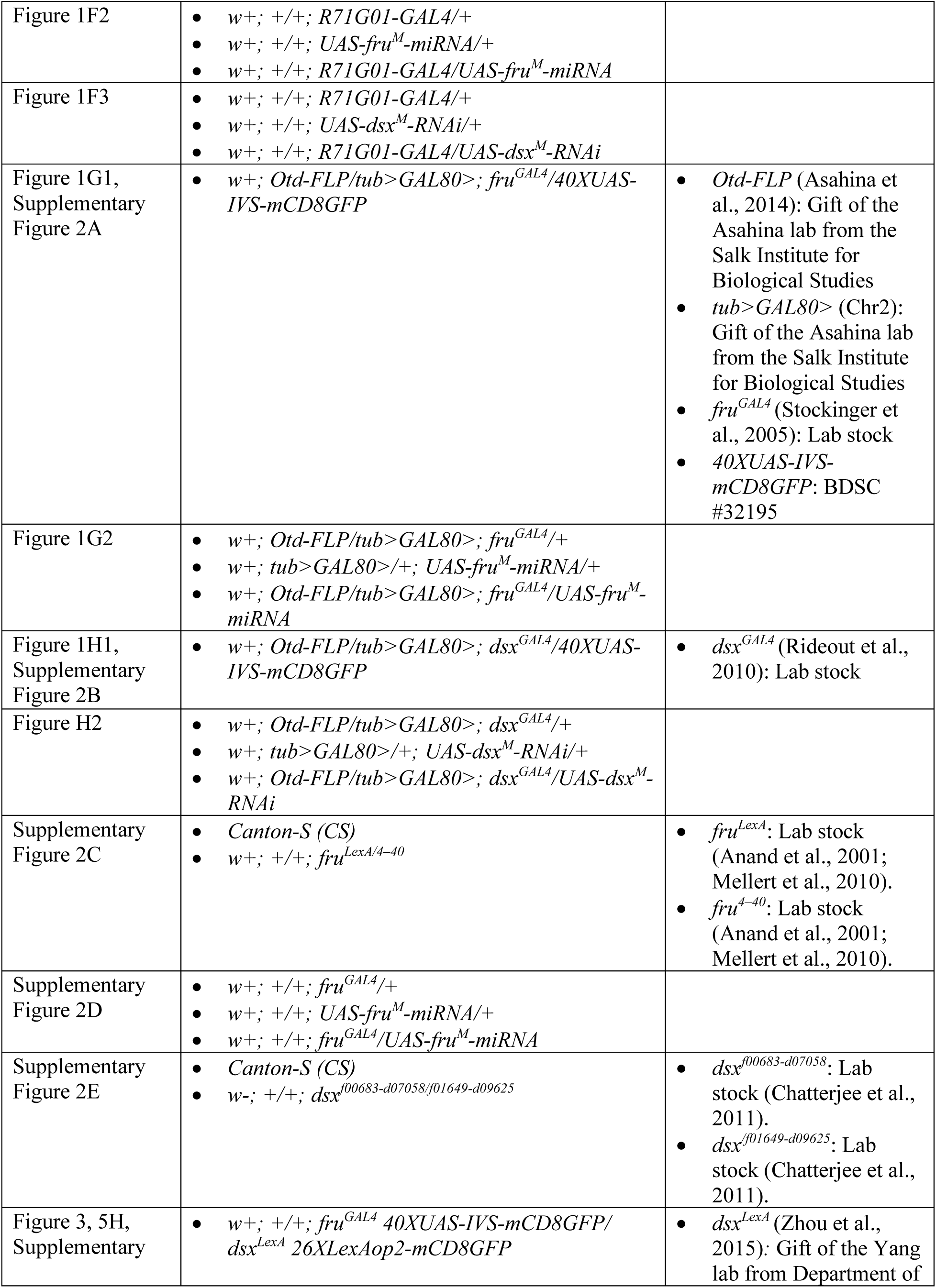
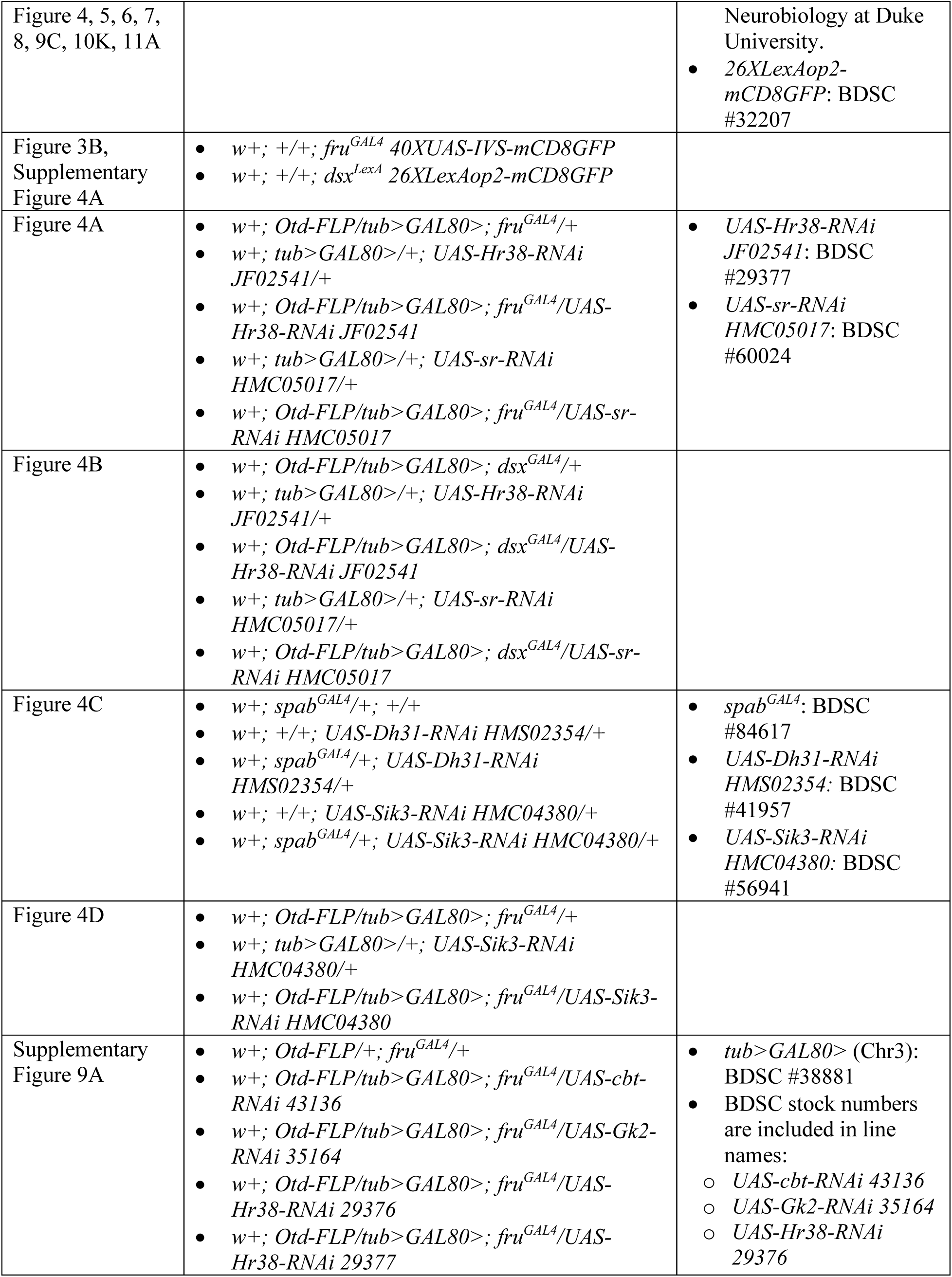
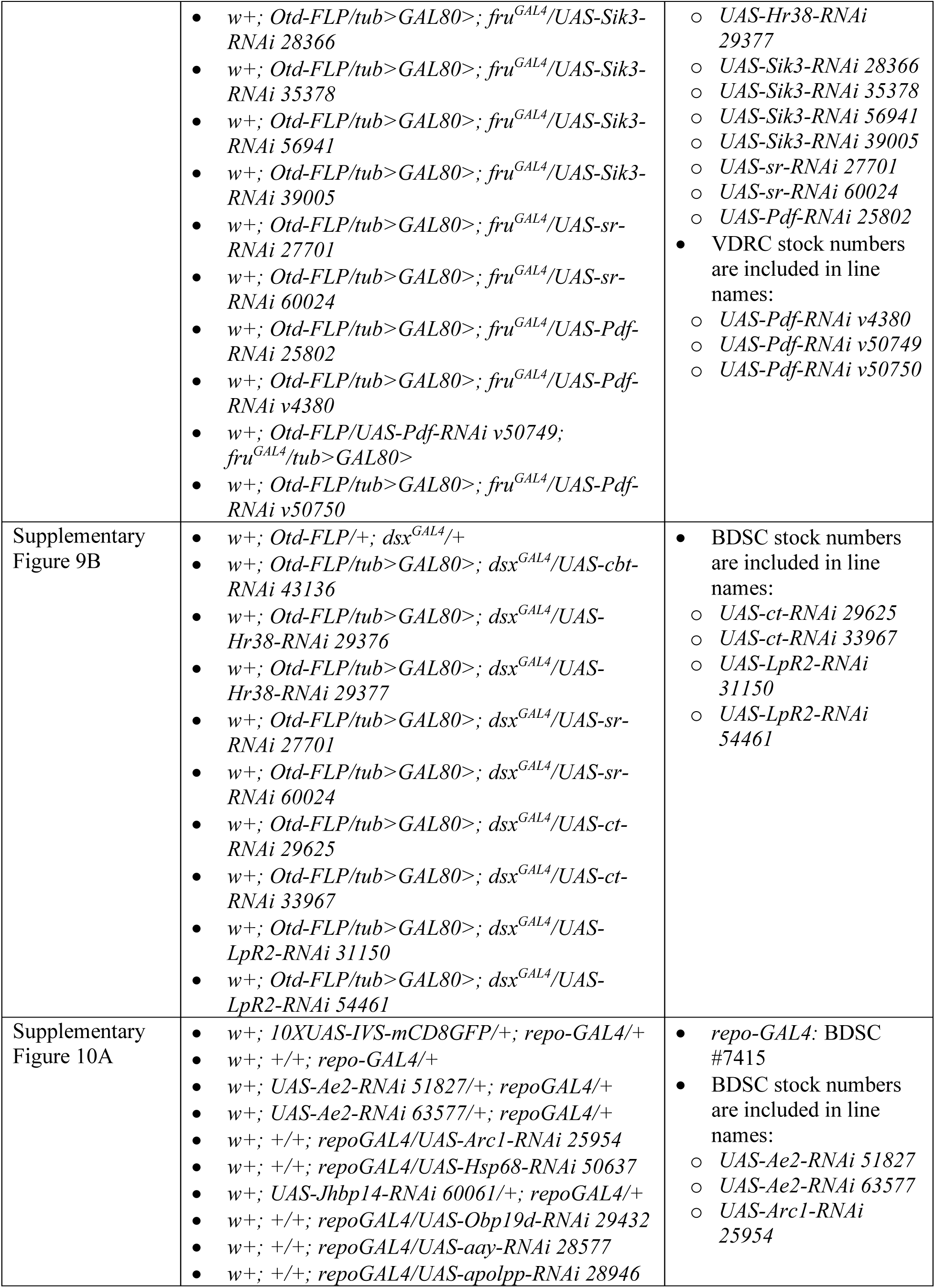
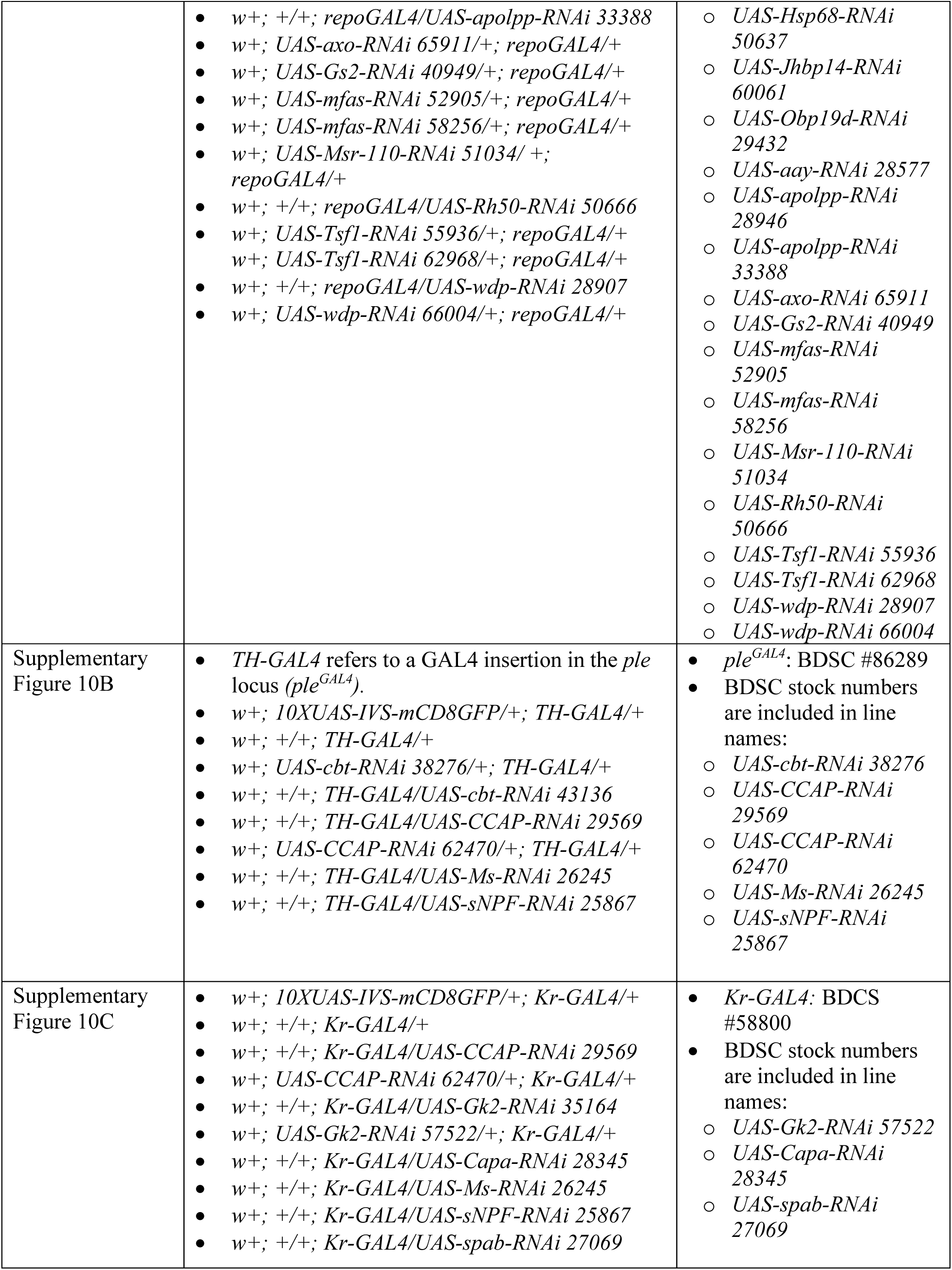
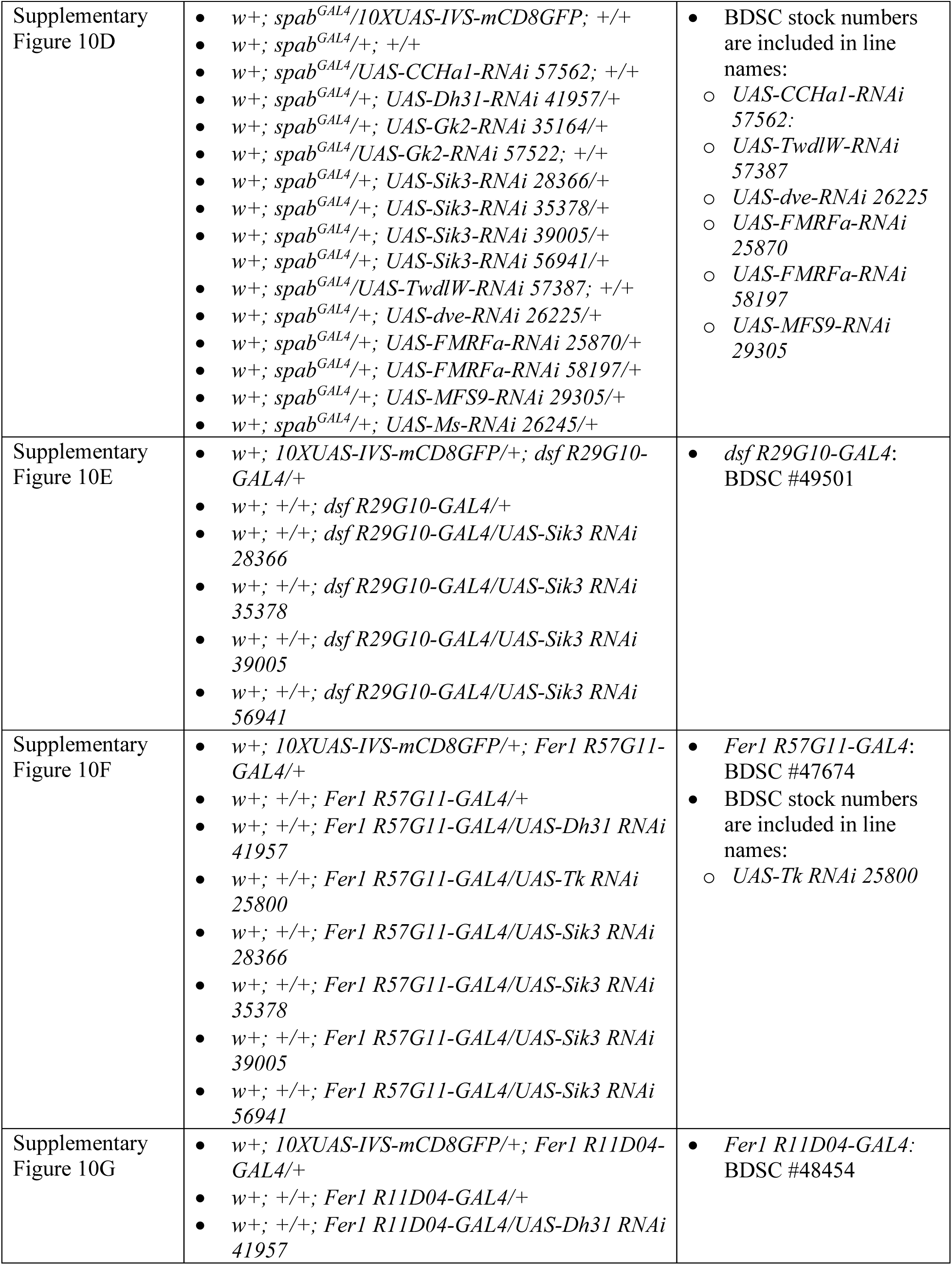
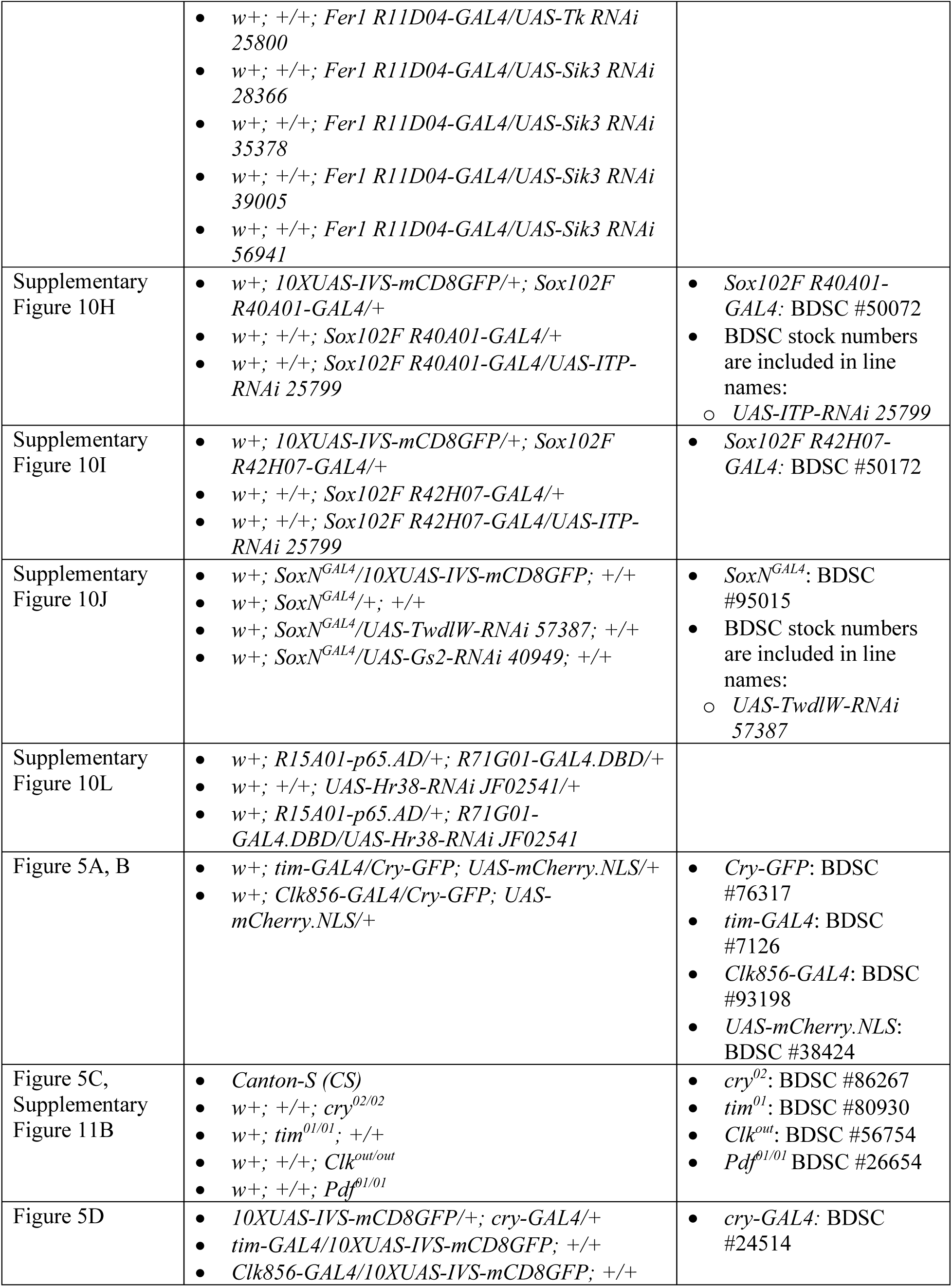
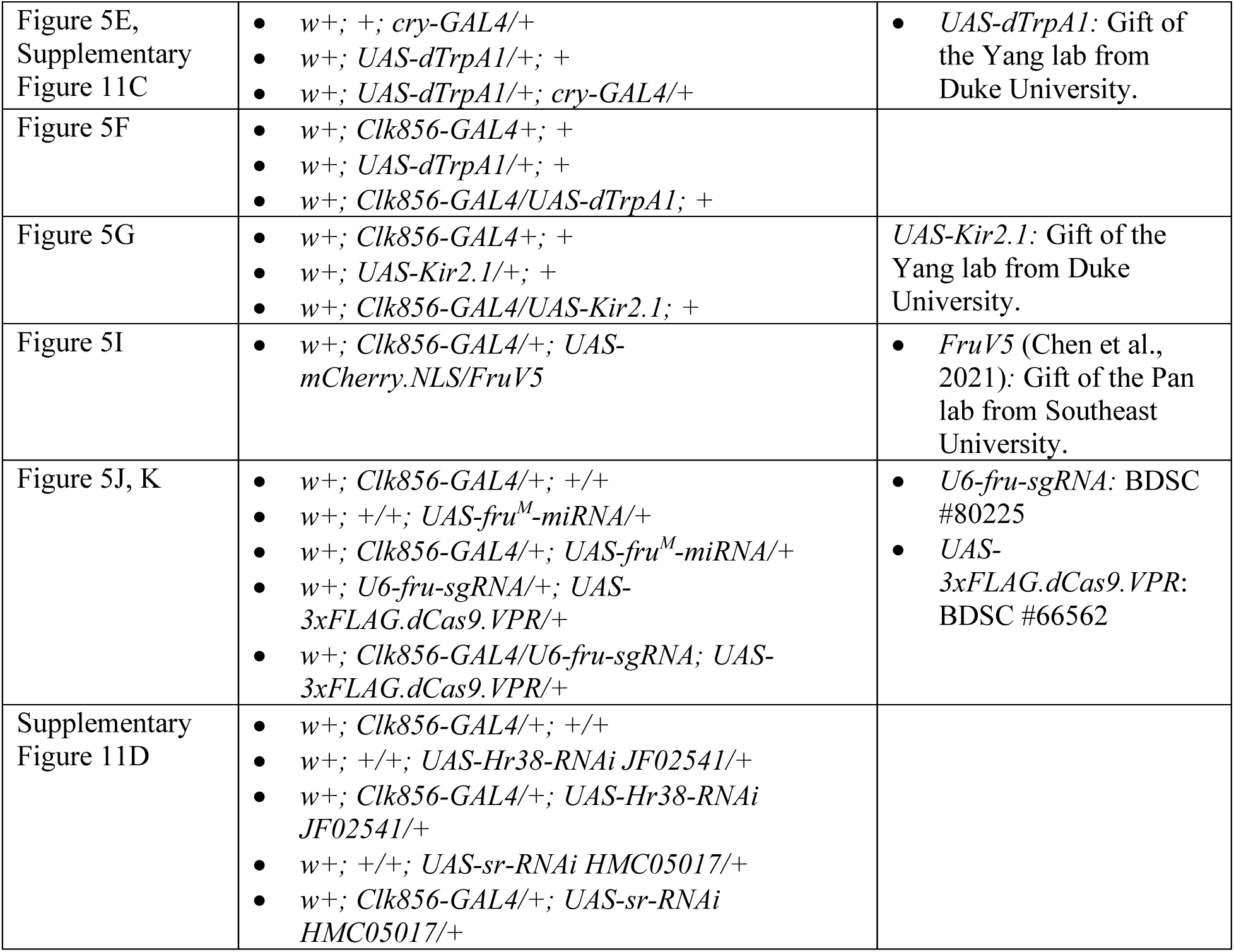
Genotypes and sources.

### Social isolation and group housing setup

Dark pupae (80–100 hours old) were picked out and separated by sex. For social isolation (single housing, SH) condition, each pupa was placed into individual vials, allowed to eclose alone and aged to 7 days. For the group housing (GH) condition, more than 30 pupae with the same sex were collected and placed into food vials, and 30 newly eclosed and unmated flies with correct genotypes were collected and put in a fresh new vial the next day of the collection day (day 0). All *w^1118^* virgin females used for courtship assay were raised under GH condition.

### Male-female single-pair courtship behavior assay setup and analysis

Courtship behavior assays were performed during ZT 7 – ZT 10. Two types of chambers were used for courtship assays. The plastic chambers were used for manual scoring blindly. Agarose chambers were designed for automated scoring to improve the efficiency of scoring courtship behavior assays.

#### Plastic chamber and manual annotation

Each plastic chamber has two layers and 10 arenas, with each arena measuring 10 mm in diameter and 2.5 mm in height in each layer. To set up the experiment, one unmated male and one virgin *w^1118^* female were gently aspirated to each arena in separate layers. When all the flies were loaded, the chamber was placed under a camera. Two layers were rotated to allow two flies to meet each other in one connected arena, and video recording was started immediately. After the recording, videos were processed by SLEAP (Pereira et al., 2022), and each arena was cropped out separately. Copulation behavior and other courtship behaviors (including chasing, orientation, tapping, single wing extension, licking, and attempted copulation) of flies in each arena were manually annotated using the Boris software (Friard and Gamba, 2016).

#### Agarose chamber and automated annotation

Each agarose chamber has 12 round arenas, with each arena measuring 25 mm in diameter and 3mm in height. To set up the experiment, one unmated male and one virgin *w^1118^* female were gently aspirated to each arena of an unused agarose chamber, separated by metal barriers before the start of the recording. When all flies were loaded, the chamber was placed at a fixed position under a camera with stable consistent light source. Barries were removed to allow two flies to meet each other, and video recording was started immediately. The trajectory of each fly was tracked by FlyTracker (Eyjolfsdottir et al., 2014) on MATLAB. Fly identity and the start time of copulation were determined manually. JAABA was trained to automatically annotate other courtship behaviors (Kabra et al., 2013).

#### Quantification

For each courtship behavior video, only the first 10 min were used for the scoring of the courtship index (CI): CI = (the time a male fly spends on courtship within 10 min)/(10 min) The time spent on copulation was also included in the CI. The results from two kinds of chambers were consistent by multiple different experiments. The housing switch (Fig. 1A-D; Supplementary Fig. 1A) and *fru* mutant (Supplementary Fig. 2C) and *dsx* mutant (Supplementary Fig. 2E) experiments presented above were performed on plastic chambers. All other courtship behavior experiments presented above were performed on agarose chambers.

### Bulk tissue RNA sequencing

#### Brain dissection and RNA extraction

Male brains were dissected in cold RNase-free 1X PBS and transferred immediately in RNAlater (Sigma-Aldrich, R0901) on ice. Each biological replicate contained 30 male fly brains. After the dissection, dilute the RNAlater using 1X RNase-free PBS with 2 times the volume of the RNAlater. Centrifuge for 1 min at max speed and remove the supernatant. Brain samples were washed with 1 ml 1X RNase-free PBS and centrifuged for 1 min at max speed. The supernatant was removed and replaced with 300 ul of TRIzol reagent (Invitrogen, 15596026). Briefly spin down to make sure all tissues are in TRIzol. Brain samples were either processed immediately or stored at –80°C for long term. To extract RNA, samples were ground sufficiently and processed with QIAGEN Shredder kit (QIAGEN, 79654) and then RNeasy Mini Kit (QIAGEN, 74104). DNase I (TURBO DNA-free Kit, Invitrogen, Thermo Fisher Scientific AM1907) was applied to remove genome DNA after the RNA extraction.

#### Library preparation and sequencing

Around 500 ng total RNA in each sample was prepared for library preparation. KAPA Stranded mRNA-Seq Kit was used to generate mRNA libraries. Libraries were sequenced on the NextSeq 2000 sequencer, generating 75-bp or 100-bp paired-end reads.

#### Bulk RNA sequencing data analysis

Raw fastq files were processed with the standard nf-core/rnaseq pipeline (Ewels et al., 2020). Briefly, reads were mapped to the *Drosophila melanogaster* BDGP6.46 genome with STAR, and read counts were generated using the Salmon method. The following analysis was performed in R (R Core Team, 2013; Wickham and Sievert, 2009). Differential expression analysis was performed with DESeq2 (Love et al., 2014). Gene ontology (GO) analyses was performed using the clusterProfiler library (Yu et al., 2012).

### Single-cell RNA sequencing

#### Single-cell suspension preparation and GFP positive cells sorting

Male fly brains were dissected in dissection buffer (Schneider′s Insect Medium, Sigma-Aldrich S0146 with 1% BSA) and transferred into dissection buffer in 1.5 ml tubes on ice. The old dissection buffer was replaced with the fresh dissection buffer. Collagenase Type I (Gibco, 17018029) stock solution was added directly into samples for a final 2mg/mL concentration. Samples were incubated at 37 °C for 20 minutes. During the incubation, samples were dissociated by gentle pipetting 20-30 times every 7 min with P200. The collagenase solution was removed after centrifugation at 300 xg for 5 min at 4°C. Cells were resuspended with 200 ul cold elution buffer (1X PBS, Gibco, 70011-044 with 0.1% BSA). Cell suspensions were passed through 30 um cell strainer caps into 5 ml collection tubes on ice. Cell strainer caps were rinsed with 800 ul cold elution buffer. Cells were processed by flow cytometry (Sony SH800 Cell Sorter) with wild-type *CS* brain cells as the negative control. GFP-positive single cells were sorted into 1.5 ml tubes and immediately transferred on ice to Duke Molecular Genomics Core (MGC) for 10X single-cell RNA seq library preparation and sequencing. Each biological replicate used thirty male fly brains, and each housing condition has two independent biological replicates. Each batch includes one GH and one SH sample processed simultaneously.

#### 10X Genomics library preparation and sequence

Cell viability and concentration were assessed by Cellometer K2 (Revvity PN CMT-K2-MX-150). Single-cell libraries were generated using Chromium X (PN 1000326), Chromium Controller (PN 110203), and 10x Genomics v3 Next GEM 3’ Single Cell chemistry (PN 1000269) following the manufacturer’s protocol, with around 20K single cells in each sample. Libraries were sequenced by the NovaSeq sequencer, and 36K –50K reads/cell were sequenced. For quality control, cDNA QCs were performed using Agilent High Sensitivity D5000 ScreenTape (Cat #5067-5592) with High Sensitivity D5000 Reagents (Cat #5067-5593). Final library QCs were performed using the Agilent High Sensitivity D1000 ScreenTape (Cat #5067-5584) with High Sensitivity D1000 Reagents (Cat #5067-5585).

#### 10X data processing

The basic analysis for the single-cell RNA sequencing followed the recommended protocols from 10X Genomics. Briefly, Cell Ranger v7.1.0 was used to demultiplex raw FASTQ files, align the reads to FlyBase release dmel-all-chromosome-r6.50 reference genome (Tweedie et al., 2009), and generate gene expression matrices for all single cells in each sample. Around 10K cells were recovered in each sample. Seurat objectives were generated for the following analysis (Satija et al., 2015). Doublets were removed by scDblFinder (Germain et al., 2022). Low-quality cells were removed (nUMI lower than 1000 or higher than 20000, gene numbers lower than 200 or higher than 4500, mitochondria genes higher than 15 percent). To do the clustering, two replicates of each housing condition were integrated first, and four datasets were merged. Clustering was performed using a standard Seurat pipeline with top 40 PCAs. Under resolution 1, the *fru*+ or *dsx*+ cells were sorted into 83 clusters in UMAPs (McInnes et al., 2018). Cell types were annotated manually based on existing cell markers (Davie et al., 2018; Park et al., 2022). Pseudo-bulk DEGs analysis were performed using DESeq2 (Love et al., 2014). Gene ontology (GO) analyses was performed using the clusterProfiler library (Yu et al., 2012).

### Immunofluorescence and confocal microscope imaging

Fly brains were dissected in cold PBT buffer (1XPBS with 10% Triton X-100) and were fixed in 4% paraformaldehyde (PFA) on nutators at room temperature for 30 minutes. Fixed brains were washed three times with fresh PBT on nutators, and every wash was 10 min at room temperature. Brain samples were stained with primary antibodies on nutators at 4 °C overnight and washed three times with fresh PBT on nutators, and every wash was 20 min at room temperature. Then, brains were stained with secondary antibodies on nutators at 4 °C overnight and washed three times with fresh PBT on nutators, and every wash was 20 min at room temperature. Antibodies used are: Rat-anti-NCad (monoclonal, DSHB-Ex#8-s, 1:20), Rabbit-anti-GFP (polyclonal, Invitrogen A11122, 1:1000), Mouse-anti-V5 (Bio-Rad, MCA1360GA, 1:200), Goat-anti-rat 647 (polyclonal, Invitrogen A21247, 1:200), Goat-anti-rabbit 488 (polyclonal, Invitrogen A11008, 1:1000), Goat-anti-mouse 488 (polyclonal, Invitrogen A11029, 1:200). Brains were mounted using Fluoromount-G Slide Mounting Medium (SouthernBiotech). Images were taken by Olympus FluoView FV1000 confocal microscope.

Fluorescence intensity quantification was performed using Python. Background-subtracted fluorescence signals were obtained by subtracting the background from the raw fluorescence intensity in each image layer, and total intensity was calculated by summing across all layers. For quantification of FruV5 signal in *Clk*-positive cells, the overlapping area between FruV5 and *Clk* signals was defined first, and FruV5 intensity within this region was calculated as described above. Statistical analyses were performed in R.

### Two-photon calcium live imaging

The two-photon calcium live imaging was performed as described previously by the Yapici lab (Mabuchi et al., 2023). Briefly, male flies were mounted on dissection holders and put in dark for one hour before the experiment. Right before imagining, the cuticle on the posterior of the head between the eyes was removed in saline. Flies were then mounted on the two-photon microscope under the objective. The P1 area was located under the scope. During the living imaging, a virgin female was delivered to a reachable position by fly forelegs to stimulate the test male fly. Neuronal responses reflected by GCaMP7s signals were recorded by the two-photon microscopy.

### Statistics

Control and experimental flies were raised and tested in parallel under identical conditions. Statistical analyses were performed using R or GraphPad. *P < 0.05, **P < 0.01, ***P < 0.001, ****P < 0.0001. All experiments were repeated in different batches. Specific animal numbers were shown in each figure or in each method part.

## Acknowledgements

We would like to thank members of the Volkan lab for discussions and help with the manuscript. We thank Dr. Yun Ding (University of Pennsylvania), Dr. Yufeng Pan (Southeast University), Dr. Kenta Asahina (Salk Institute for Biological Studies) and Dr. Rebecca Chung-Hui Yang (Duke University) for kindly sharing fly lines. We would like to acknowledge the Duke Molecular Physiology Institute Molecular Genomics Core, especially Karen Abramson, Ellora Haukenfrers and Michael Aksu for their help on the generation of single cell RNA sequencing data. We thank the Duke High Throughput Sequencing Facility, Bloomington Stock Center, Developmental Studies Hybridoma Bank, Vienna Drosophila Resource Center, and Drosophila Genomics Resource Center for their services.

## Author contributions

Conceptualization: CD and PCV. Investigation: CD, SO, LJ, MRB and SS with help from YM, SA, LG, SR, SK, EB, and JL. Analysis/interpretation of data: CD, JESF, CDJ, NY, and PCV. Manuscript— original draft: CD and PCV.

## Figures

**Supplementary Figure 1.**
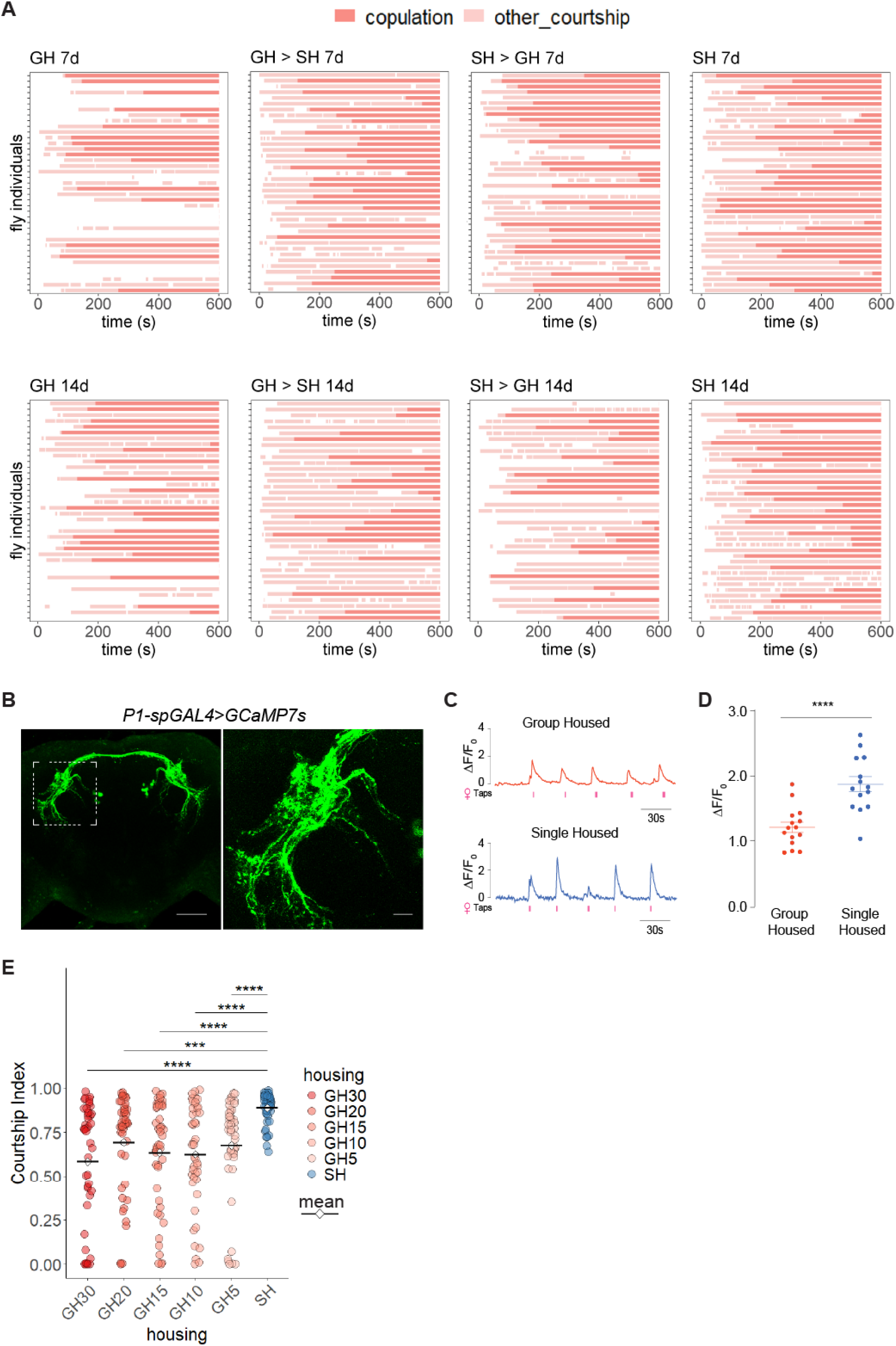
Courtship behavior assays and P1 neuron live imaging in male flies. **(A)** Ethograms showing copulation and other courtship behaviors for each individual fly raised under four different housing conditions. **(B)** P1 neurons labelled by *P1-spGAL4>GCaMP7s*, stained for GFP. Live imaging location used in **(C-D)** is indicated by white, dotted box. **(C)** Representative traces of P1 evoked responses in GH and SH males in response to the taste of females (red ticks indicate time of foreleg taps). **(D)** Statistics of average peak response (ΔF/F_0_) for P1 neurons evoked by female stimulation. Data points represent individual males. Unpaired t-test was applied. Significance of P-value: ****P < 0.0001. **(E)** Courtship Index in male-female single pair assays with male flies raised in different grouping numbers. The Kruskal–Wallis rank sum test followed by Dunn’s multiple comparisons test was applied. Significance of adjusted P-value (P_adj): *P_adj < 0.05, **P_adj < 0.01, ***P_adj < 0.001, ****P_adj < 0.0001.

**Supplementary Figure 2.**
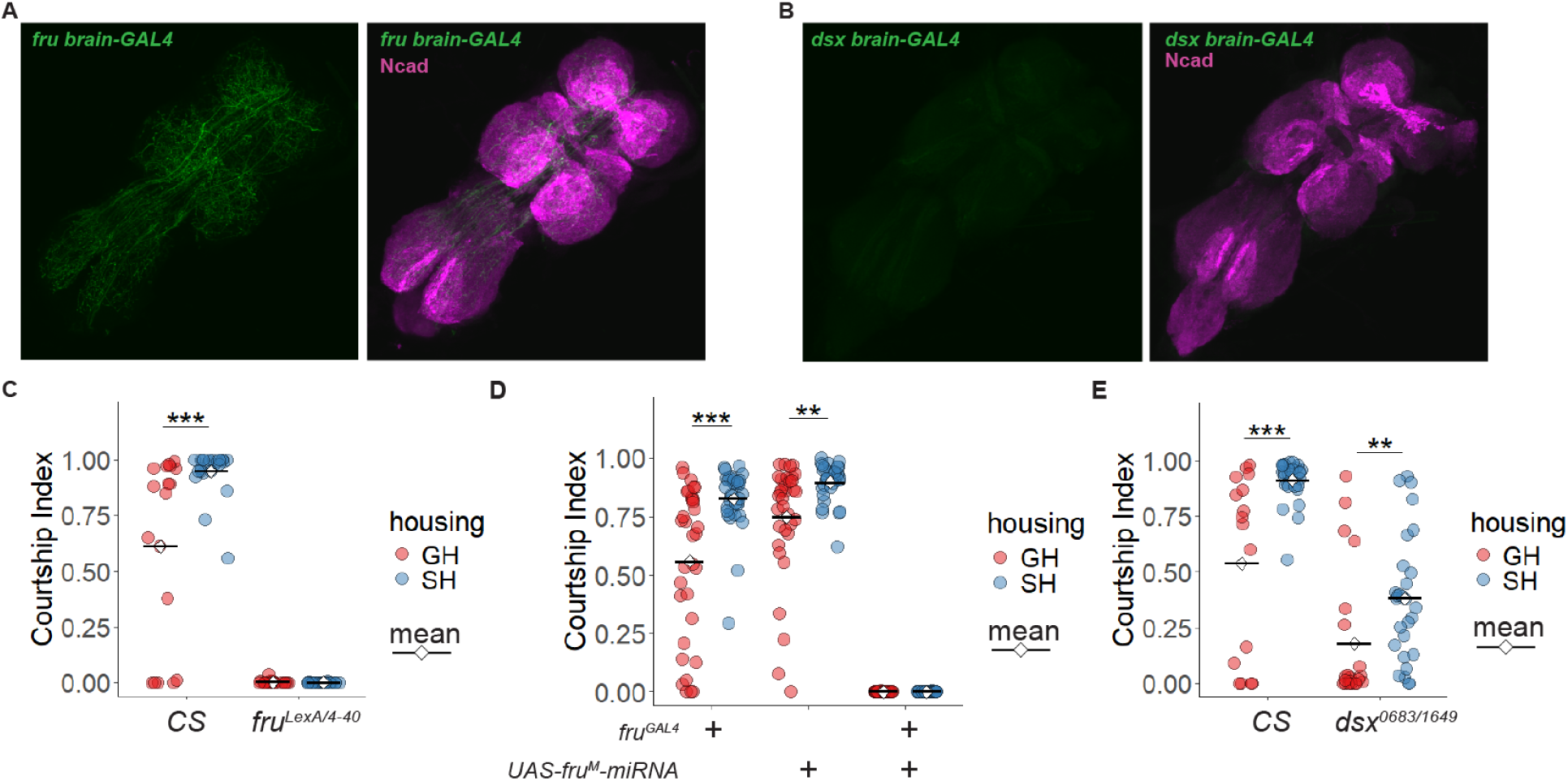
The effect of *fru* and *dsx* on male courtship behavior. **(A)** Expression pattern of *fru brain-GAL4> mCD8::GFP* is absent from the ventral nerve cord, stained for GFP (green) and Ncad (magenta). **(B)** Expression pattern of *dsx brain-GAL4> mCD8::GFP* in the ventral nerve cord, stained for GFP (green) and Ncad (magenta). **(C)** Courtship Index from male–female single-pair assays with males of wild-type *CS* and *fru* mutants. **(D)** Courtship Index from male–female single-pair assays in control and experimental groups with *fru* knockdown in all *fru*-positive cells. **(E)** Courtship Index from male–female single-pair assays with males of wild-type *CS* and *dsx* mutants. The Wilcox test was applied to the comparison between GH and SH conditions of the same genotype. Significance of adjusted P-value (P_adj): *P_adj < 0.05, **P_adj < 0.01, ***P_adj < 0.001, ****P_adj < 0.0001.

**Supplementary Figure 3.**
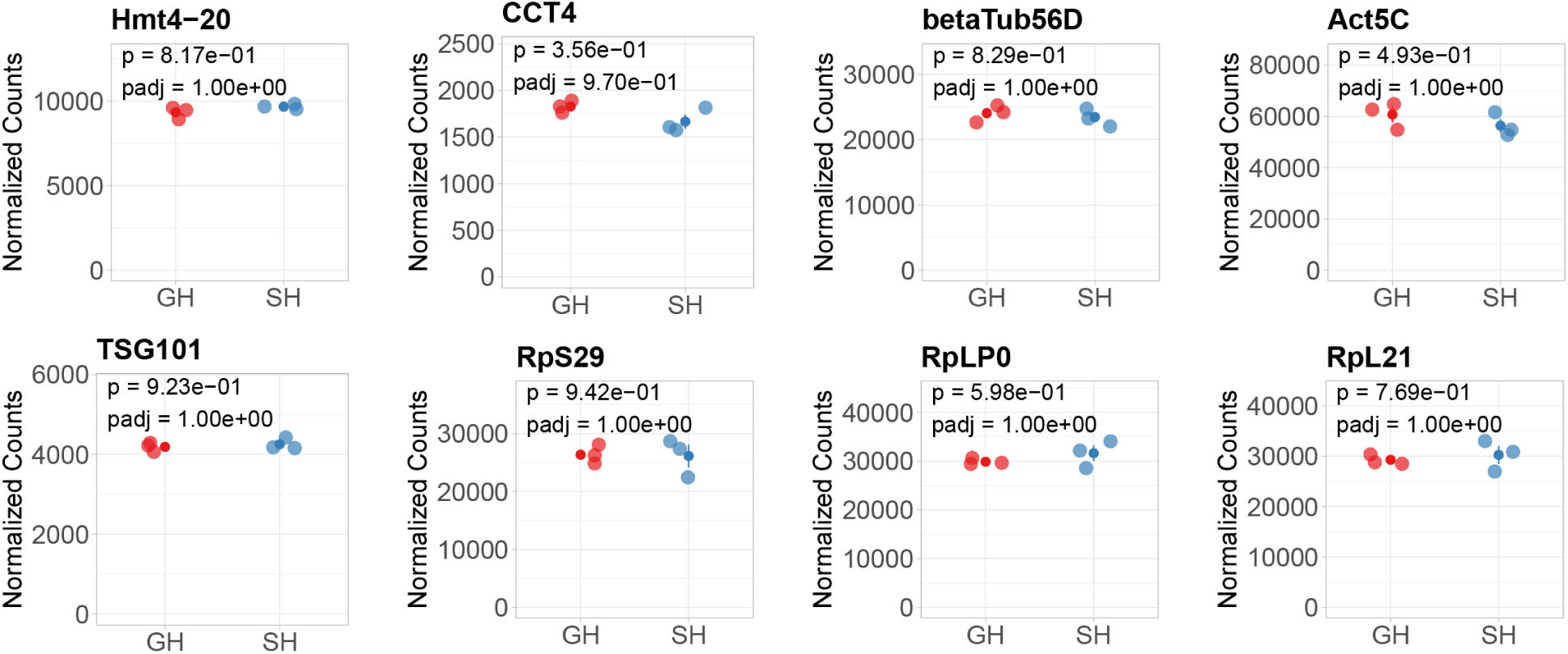
Transcript levels for representative housekeeping genes among all male brain samples in the bulk tissue RNAseq.

**Supplementary Figure 4.**
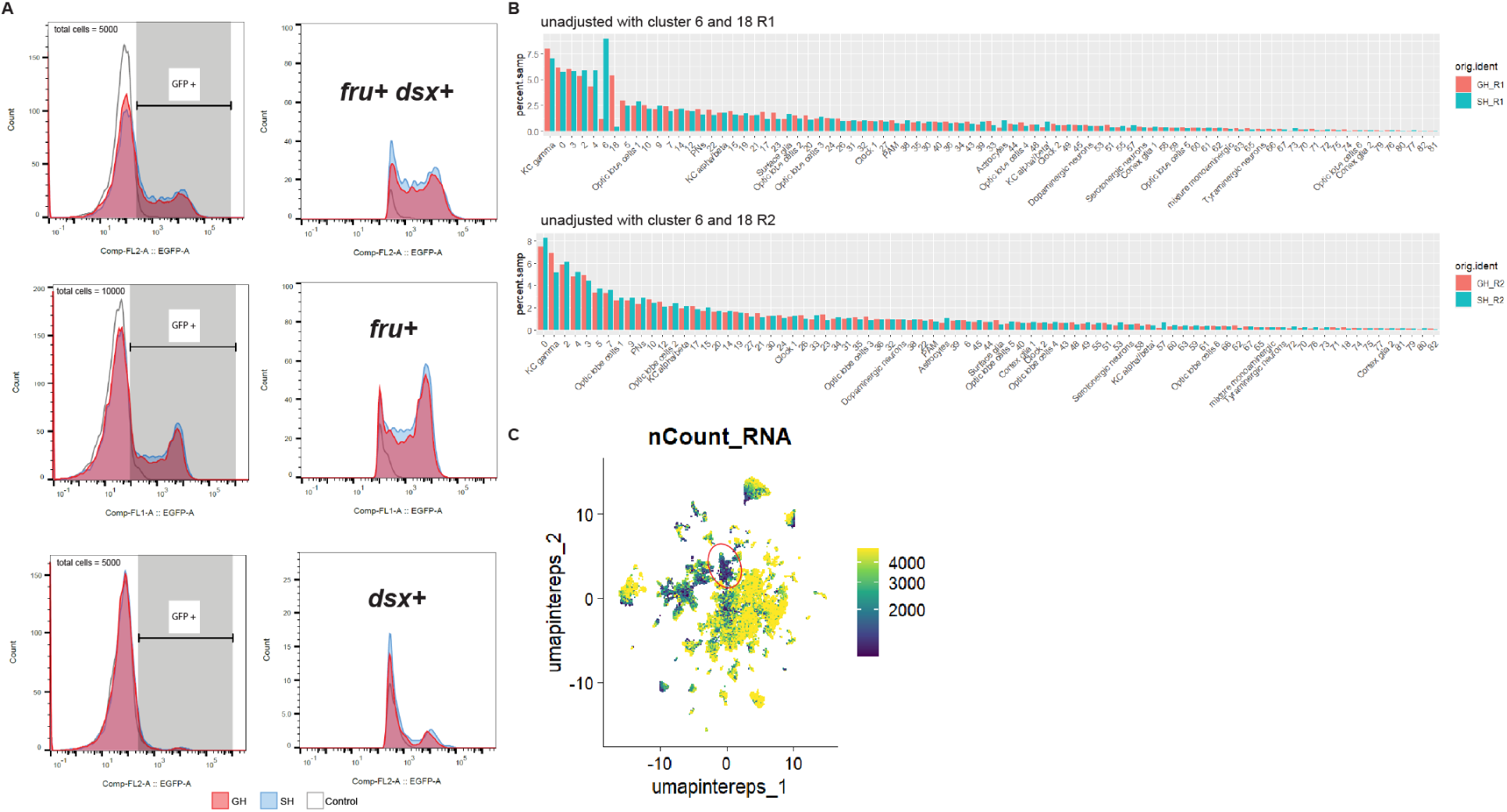
The proportions of different cell clusters in GH and SH male brains from single-cell RNAseq. **(A)** Proportion of GFP-labeled combined *fru*⁺ cells and *dsx*^+^ cells, *fru*^+^ only cells, and *dsx^+^*only cells. The X-axes indicate GFP fluorescence intensity. Gray-shaded regions (GFP⁺) are magnified in the right panels and quantified in Figure 3B. Red and blue represent GH and SH conditions, respectively; gray lines indicate negative controls from *CS* brain cells. **(B)** Comparison of cluster proportions in GH and SH conditions in two biological replicates. **(C)** UMAP plot illustrating the overall level of transcript counts.

**Supplementary Figure 5.**
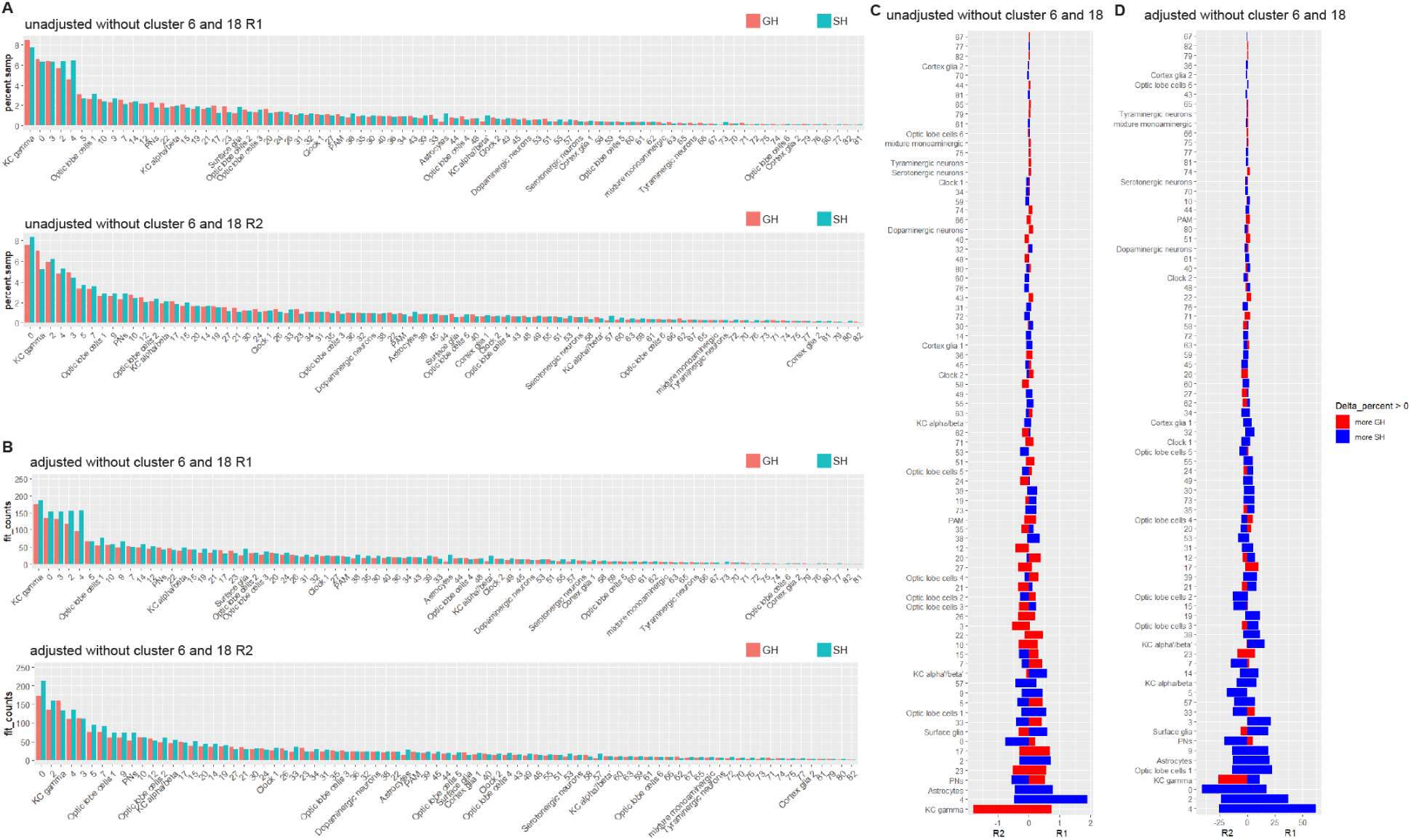
The increase in the number of *fru-* and *dsx-*positive cells originate from many clusters in the SH condition. **(A)** Comparison of cluster proportions in GH and SH conditions in two biological replicates, excluding clusters 6 and 18. **(B)** Comparison of processed counts in GH and SH conditions, adjusted for the actual percentage of GFP⁺ cells measured by FACS, with clusters 6 and 18 excluded. **(C)** Differences in cluster proportions between GH and SH conditions across two biological replicates in (A). Red indicates clusters with a higher proportion in GH, while blue indicates clusters with a higher proportion in SH. **(D)** Adjusted differences in cluster counts between GH and SH conditions across two biological replicates in (B). Red indicates clusters with more cells in GH, while blue indicates clusters with more cells in SH.

**Supplementary Figure 6.**
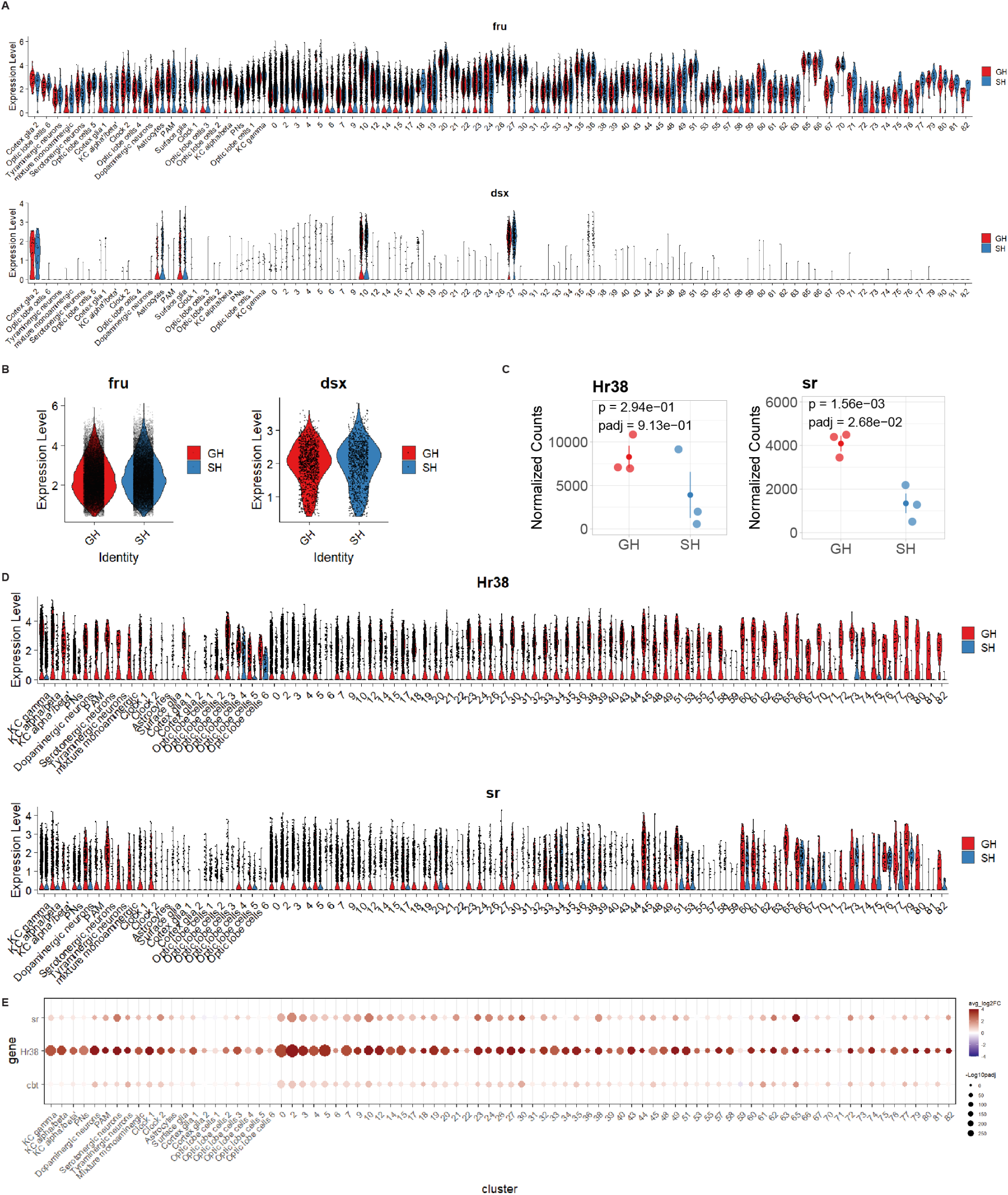
The expression pattern of *fru, dsx, Hr38,* and *sr* in each cell cluster in single-cell RNA sequencing. **(A)** Violin plots comparing the distribution of *fru* and *dsx* expression levels across different cell clusters in GH and SH conditions in single-cell RNA sequencing. **(B)** Violin plots comparing the distribution of overall *fru* and *dsx* expression levels in cells with detectable *fru* level or *dsx* level, respectively, in GH and SH conditions in single-cell RNA sequencing. **(C)** Transcript levels for *Hr38* and *sr* among all male brain samples in the bulk tissue RNAseq. **(D)** Violin plots comparing the distribution of *Hr38* and *sr* expression levels across different cell clusters in GH and SH conditions in single-cell RNA sequencing. **(E)** Results of pseudobulk analysis showing Log2FoldChange (GH/SH) and –Log10(P-adj) of *sr, Hr38* and *cbt* in each cell cluster.

**Supplementary Figure 7.**
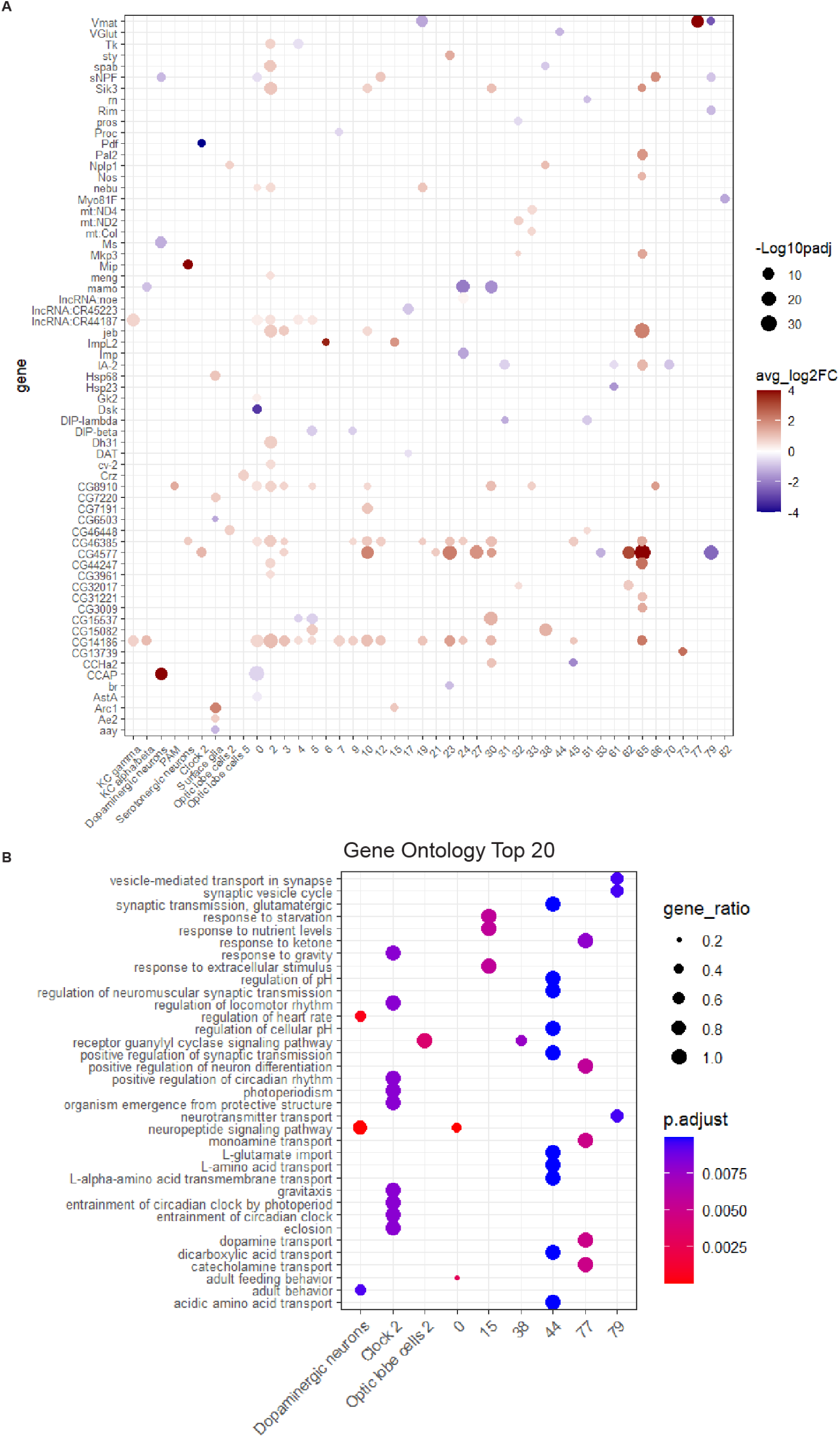
DEGs of each cell cluster from the GH vs SH comparison in single-cell RNA sequencing. **(A)** Results of pseudobulk analysis showing Log2FoldChange (GH/SH) and – Log10(P-adj) of DEGs in each cell cluster. **(B)** Top 20 most significantly enriched GO terms for DEGs of GH/SH comparisons in different cell clusters from (A) (q-value < 0.01).

**Supplementary Figure 8.**
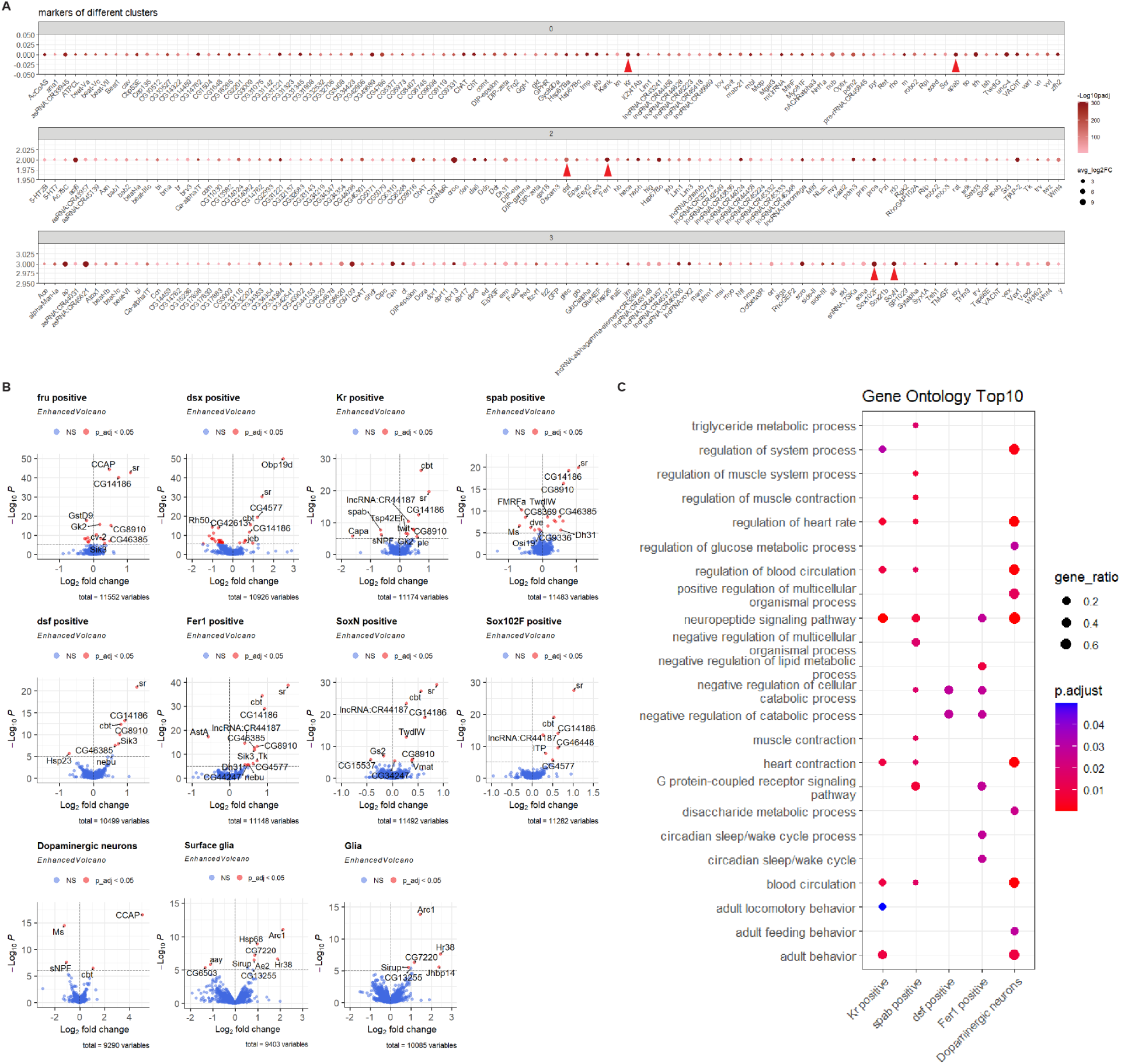
DEGs in different subsets defined by marker genes in single-cell RNA sequencing. **(A)** Marker genes for cell cluster 0, cluster 2, and cluster 3 from the UMAP clusters in Figure 3D. **(B)** Volcano plots displaying Log2FoldChange (GH/SH) and –Log10(P-value) of genes in different subsets defined by marker genes. Red indicates DEGs from pseudobulk analysis. **(C)** Top 10 most significantly enriched GO terms for DEGs of GH/SH comparisons in different cell subsets from Figure 3F (q-value < 0.05).

**Supplementary Figure 9.**
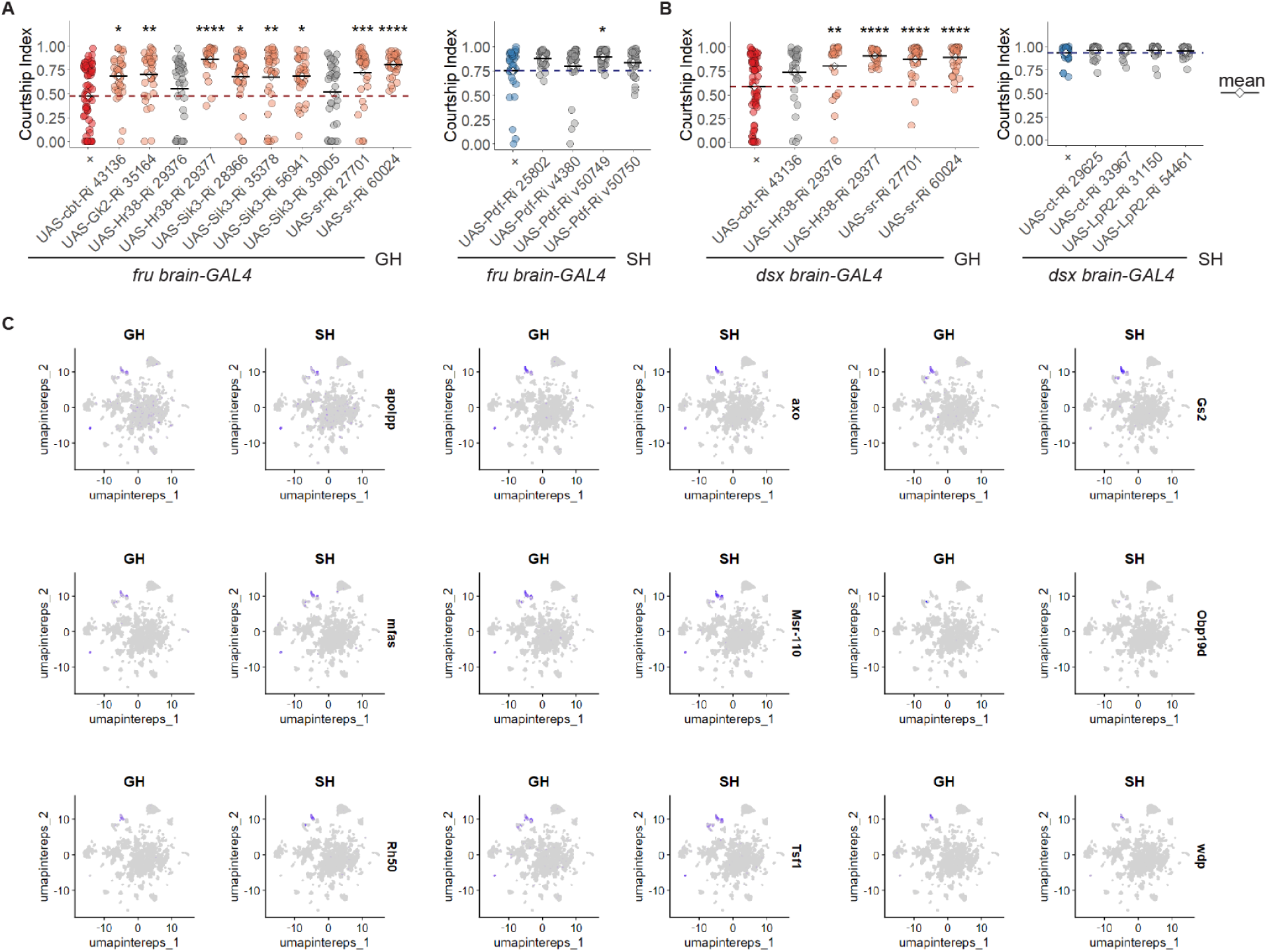
RNAi screening of DEGs in *fru* or *dsx* brain neurons. (**A-B**) Courtship index of male flies with RNAi-mediated knockdown of selected DEGs in *fru-* **(A)** or *dsx*-**(B)** expressing brain neurons. In each plot, the first column (+) represents the control group, with red indicating GH conditions and blue indicating SH conditions. The dashed lines indicate the mean courtship index for control groups. Each dot represents an individual male fly, with light red dots indicating that the knockdown group exhibited significantly higher courtship levels compared to the control, and light blue dots indicating that the knockdown group exhibited significantly lower courtship levels compared to the control. The Wilcox test was applied to the comparison between each RNAi group and the control group. Significance of adjusted P-value (P_adj): *P_adj < 0.05, **P_adj < 0.01, ***P_adj < 0.001, ****P_adj < 0.0001. **(C)** Visualization of the expression pattern of selected genes. Each dot is one cell colored by the expression levels of each gene.

**Supplementary Figure 10.**
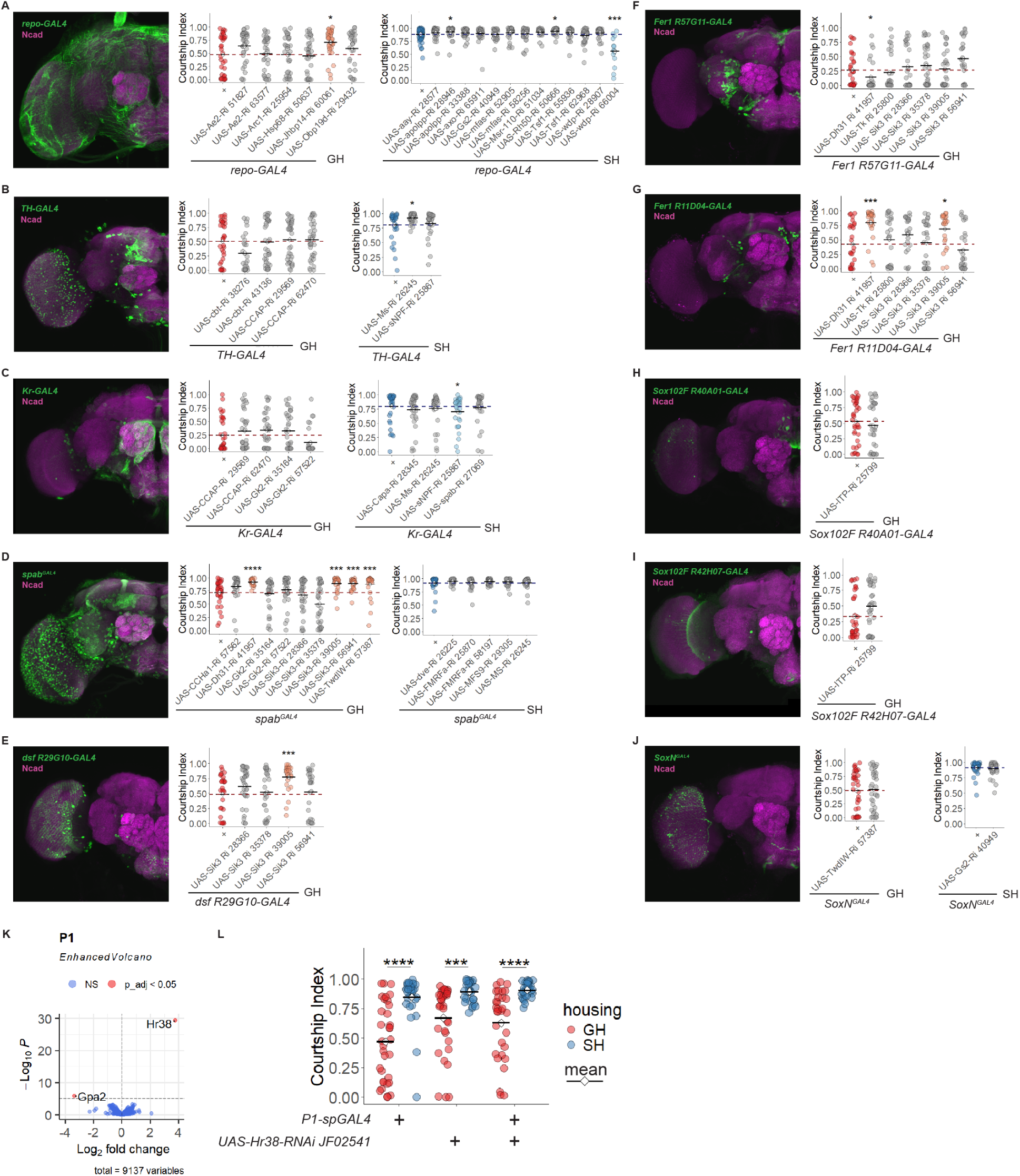
RNAi screening identifies candidate genes mediating social experience-dependent courtship plasticity. (**A-J**) RNAi-mediated knockdown screening of selected DEGs in *repo-* **(A)**, *ple*-(TH) **(B)**, *Kr-* **(C)**, *spab-* **(D)**, *dsf-* **(E)**, *Fer1-* **(F-G),** *Sox102F-* **(H-I)**, and *SoxN-* **(J)** expressing cells. Left: Cells labelled by different drivers in the brain, stained for GFP (green) and Ncad (magenta). Right: Courtship index of male-female courtship assays in each screening group. In each plot, the first column (+) represents the control group, with red indicating GH conditions and blue indicating SH conditions. The dashed lines indicate the mean courtship index for control groups. Each dot represents an individual male fly, with light red dots indicating that the knockdown group exhibited significantly higher courtship levels compared to the control, and light blue dots indicating that the knockdown group exhibited significantly lower courtship levels compared to the control. The Wilcox test was applied to the comparison between each RNAi group and the control group. Significance of adjusted P-value (P_adj): *P_adj < 0.05, **P_adj < 0.01, ***P_adj < 0.001, ****P_adj < 0.0001. **(K)** Volcano plots displaying Log2FoldChange (GH/SH) and –Log10(P-value) of genes in P1 neurons. Red indicates DEGs from pseudobulk analysis. **(L)** Courtship Index from male–female single-pair assays in control and experimental groups with *Hr38* knockdown in P1 neurons with the P1-spGAL4 driver. The Wilcox test was applied to the comparison between GH and SH conditions of the same genotype. Significance of adjusted P-value (P_adj): *P_adj < 0.05, **P_adj < 0.01, ***P_adj < 0.001, ****P_adj < 0.0001.

**Supplementary Figure 11.**
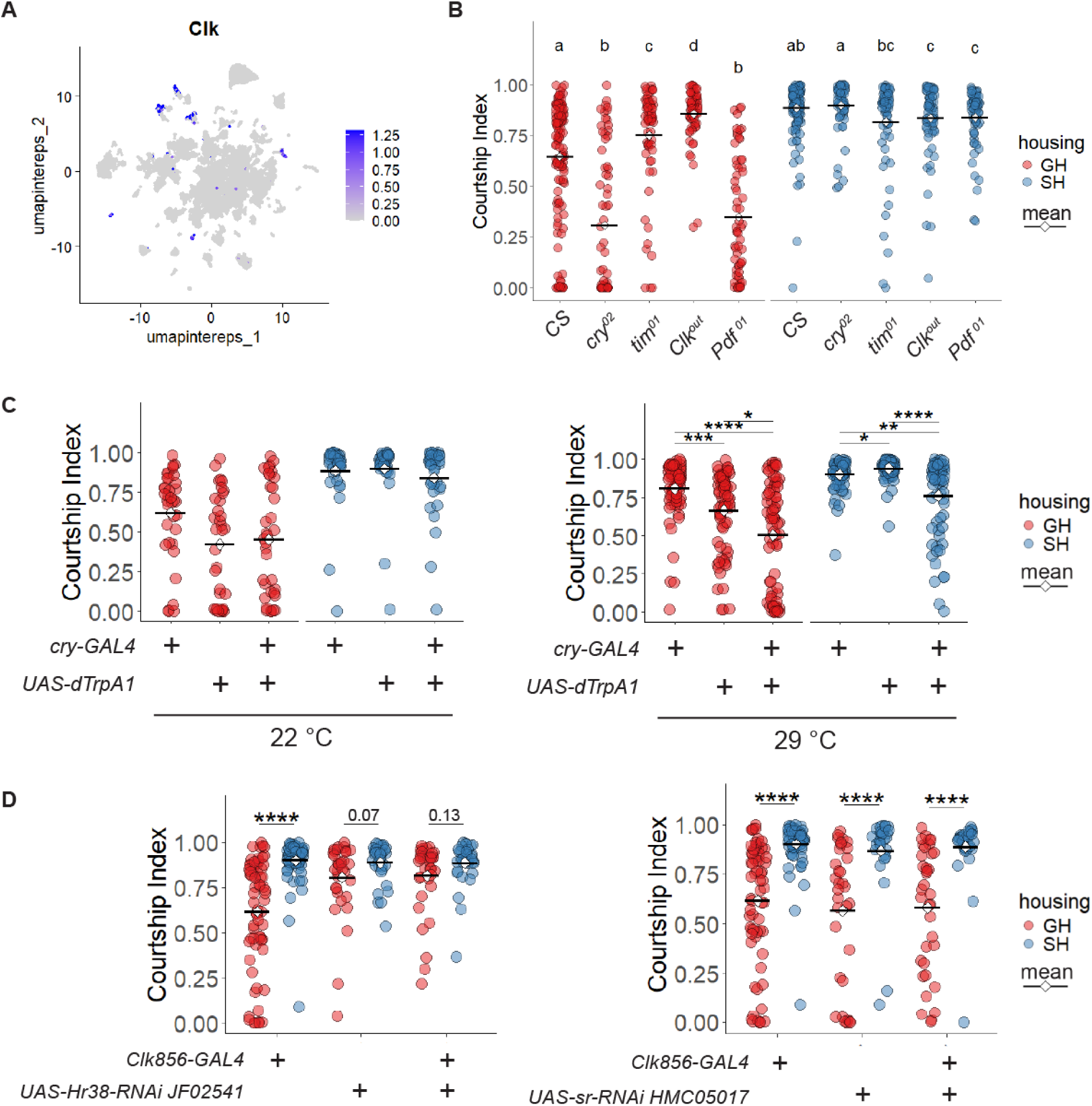
The effect of the clock system and immediate early genes in social experience-dependent modulation of courtship. **(A)** Visualization of the expression pattern of *Clk* in the single-cell RNAseq. **(B)** Courtship Index from male–female single-pair assays in *CS*, and mutants in *cry*, *tim*, *Clk* and *Pdf*. **(C)** Courtship Index from male–female single-pair courtship assays with *cry-GAL4* driving dTrpA1 under 22°C and 29°C. **(B-C)** The Kruskal–Wallis rank sum test, followed by Dunn’s multiple comparisons test, was applied for the comparison among different genotypes grouped by housing. **(D)** Courtship Index from male–female single-pair assays comparing control and experimental groups of *Hr38* and *sr* knockdown in *Clk* expressing cells. The Wilcox test was applied to the comparison between GH and SH conditions of the same genotype. Significance of adjusted P-value (P_adj): *P_adj < 0.05, **P_adj < 0.01, ***P_adj < 0.001, ****P_adj < 0.0001.

